# Temporal resolution trumps spectral resolution in UAV-based monitoring of canopy senescence dynamics

**DOI:** 10.1101/2023.12.13.571461

**Authors:** Flavian Tschurr, Lukas Roth, Nicola Storni, Olivia Zumsteg, Achim Walter, Jonas Anderegg

**Author notes:** Contributing authors.

## Abstract

Senescence is a dynamic process that is affected by many environmental, genetic, and physiological factors. Quantifying this process is important for breeding wheat varieties with high yield and of high quality. We present a method that allows up-scaling of the state of the art method - visual scoring - by using image sequences acquired from Unmanned Aerial Vehicles (UAV). This reduces measurement time and environmental changes during the measurement as well as rater bias. We compared the potential of a widely used multispectral sensor and a cheaper high-resolution RGB camera to track the dynamics of senescence. A UAV each was equipped with one of these sensors and used to measure canopy reflectance throughout the senescence process that lasted several weeks, for more than 400 winter wheat cultivars across three field seasons. Multiple spectral and RGB indices were calculated at the experimental plot level and used to model the dynamics of senescence. Model fits were further processed to extract key time points of the senescence phase. By comparing the results of the two sensors with each other and with the visual evaluation, respectively, we show that both sensors allow monitoring of senescence dynamics and measure key time points of the phase with a precision close to that of more sophisticated proximal sensing approaches. Optimal timing of measurements proved to be more important than the choice of sensor, confirming that timely and frequent measurements should be prioritized over more expensive sensors that provide a higher spectral resolution.

## 1 Background

A significant portion of daily calorie and protein intake is based on on a handful of arable crops, notably wheat. Consequently, there is an immediate need to identify and mitigate risks associated with the cultivation of these crops (Tilman et al. 2011). This urgency is further accentuated by the projected decline in global wheat production and quality, including protein content and composition, due to the impacts of global climate change (S. Asseng et al. 2015; Senthold Asseng et al. 2019). Potential mitigation strategies encompass diversifying cropping systems or developing crop varieties with enhanced performance under stress conditions (Reynolds et al. 2016). An important stress-adaptive trait that is effective in several crops is the prolonged maintenance of green leaf area after anthesis to increase carbon assimilation during grain filling. (Thomas and Smart 1993).

In wheat, the primary limiting factor for potential grain yield is seen predominantly in sink strength, which refers to the number of grains available for grain filling and their capacity to absorb assimilates. This critical determinant is largely established before grain filling initiates, extending up to and including a short period after anthesis, as extensively reviewed by Borŕas et al. 2004, R. A. Fischer 2008 and Distelfeld et al. 2014. Nevertheless, several studies have documented a positive correlation between delayed senescence and grain yield, especially in combination with stress conditions (Verma et al. 2004; LopesMaimaitijiang et al. 2012; Montazeaud et al. 2016; M. Christopher et al. 2017). In addition, the onset of senescence marks a basic transition of canopies from carbon assimilation to N remobilization (Thomas and Ougham 2014), and its timing is therefore expected to affect not only GY but also grain protein concentration, which is a key quality parameter of major economic importance.

In situations in which severe stress challenges wheat, a prolonged retention of green leaf area can be construed as a strategy to prevent premature senescence. An example of this is the CIMMYT wheat line SeriM82, which has a significant stay-green related yield advantage in Australian multi-environment trials over locally adapted genotypes (J. T. Christopher et al. 2008). In this case, a denser root system at depth likely increases deep soil moisture extraction late in the season, supporting an extended grain filling duration by delaying stress-induced senescence, resulting in increased individual grain weight (J. T. Christopher et al. 2008; John T. Christopher, Veyradier, et al. 2014; John T. Christopher, M. J. Christopher, et al. 2016). Understanding this yield-determining phase in wheat is critical, especially in light of future climate change and more extreme events (Seneviratne et al. 2021). Even in countries with temperate conditions, such as Switzerland (A. M. Fischer et al. 2022), an increase in agriculturally relevant climate extremes, such as heat waves, is expected (Tschurr, Feigenwinter, et al. 2020).

There are several approaches to track senescence in crop plants (Kipp et al. 2014; Anderegg, Yu, et al. 2020). This is often done with very expensive equipment, such as hyperspectral sensors (spectroradiometry). These sensors are used in manual plot-by-plot measurements and require optimal environmental conditions for the entire duration of the measurement, in particular stable light conditions. Therefore, there is a great need to increase the throughput, for example by using unmanned aerial vehicle (UAV)s derived traits (Makanza et al. 2018; Li et al. 2023). Although such approaches can reduce human bias compared to visual scoring and are thus more reproducible, they require stable measurement conditions. In particular, different lighting conditions can strongly influence the measurements and cause bias if they change during a measurement. Therefore, faster measurement approaches, such as using UAVs, allow a high measurement density under good conditions, as the time per measurement is lower compared to other approaches. This can additionally reduce costs, as less labor is required and very expensive sensors are not needed. Furthermore, for dynamic and complex processes such as senescence, multi-environment trials are required to disentangle genotypic and environmental effects. Therefore, it is very important that a reproducible and unbiased method is used to meaningfully compare across different environments. Furthermore, the use of less expensive sensors, such as regular Red-Green-Blue (RGB) cameras, would be beneficial to make such measurements accessible to a wide range of applications. Previous studies demonstrated that vegetation index dynamics utilizing few specific spectral bands provide an accurate and robust representation of canopy senescence dynamics in wheat, whereas approaches based on full reflectance spectra suffered from specificity to environments (Anderegg, Yu, et al. 2020; Hassan et al. 2018). This suggests that senescence dynamics may be accurately tracked using less sophisticated and readily affordable sensors. The aim of this study was therefore i) to evaluate UAV-based reflectance measurements as tools for high throughput senescence monitoring; ii) to compare the state-of-the-art approach relying on expensive multispectral (SPC) sensors with less sophisticated and more affordable RGB cameras, and with the conventional method of visual scoring; and iii) to evaluate the effect of the temporal measurement resolution in comparison to the sensor used. Four winter wheat trials, conducted across three years, each encompassing between 180 and 800 plots, were considered in this study to develop and validate a method for the above research objectives. Phenologically and morphologically diverse winter wheat cultivars were included in the investigation: The GABI wheat panel (Kollers et al. 2013; Gogna et al. 2022), European and Swiss cultivars, as well as more than 300 breeding lines were measured over the years. Accordingly, these trials were well suited for the development and evaluation of a robust method with respect to both a wide genetic diversity as well as year-to-year variability.

## 2 Materials and Methods

The analysis is divided into four main steps: data acquisition, pre-processing (photogrametry for RGB and SPC, signal correction for SPC), dynamic modelling, and validation. Data from two sensors, a SPC and a RGB-camera mounted on UAVs were combined with visual scorings. Extensive pre-processing of the UAV raw data is required to extract plot-level reflectance for each measurement time point. Within the dynamic modeling, the best index and modelling approaches are considered and used to extract key time points of the senescence phase. These time points were then used to compare and validate the two sensors with regard to their capability of capturing visually observed senescence dynamics (Figure 1).

**Fig. 1.**
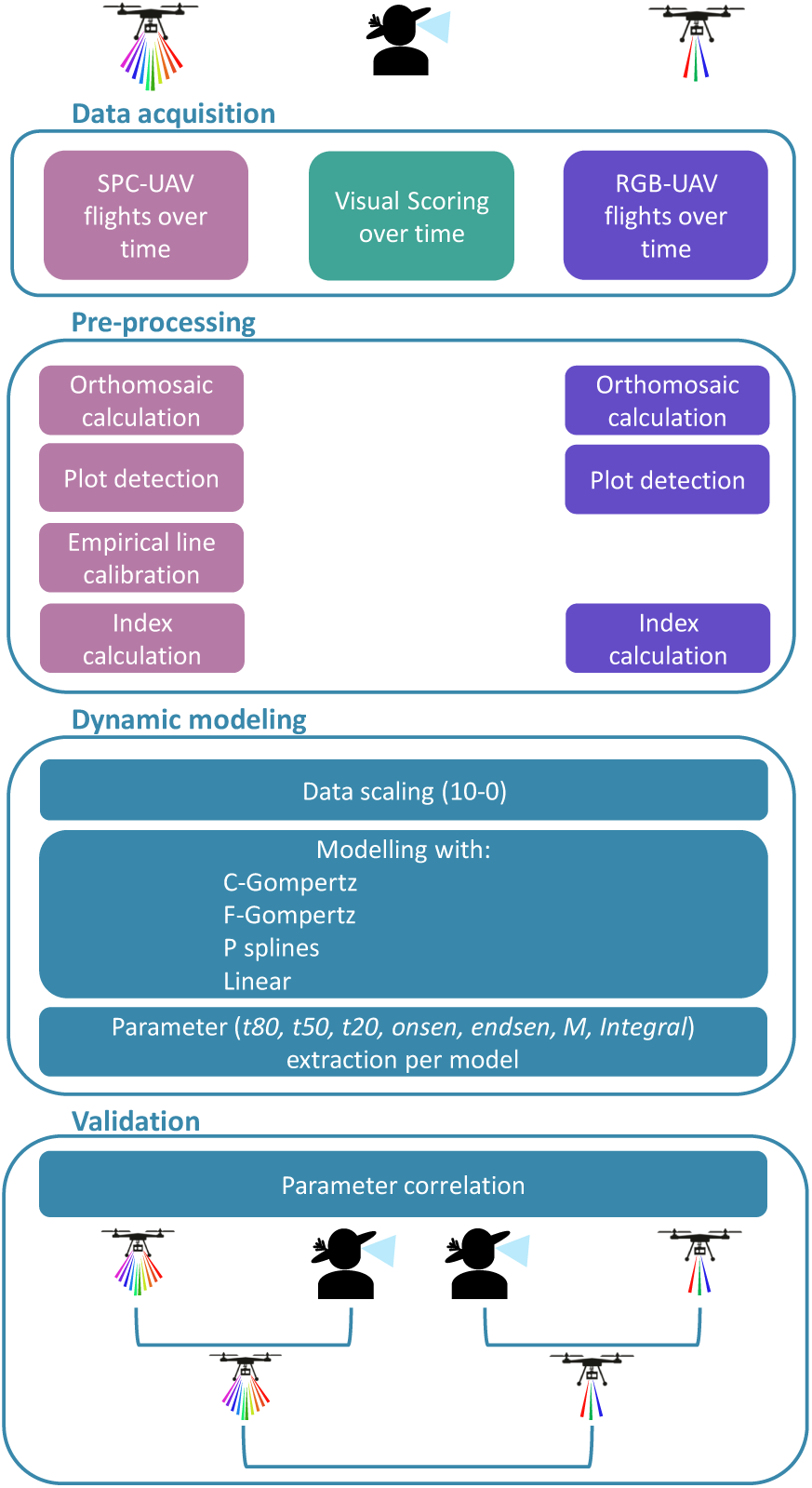
Overview of the workflow in this paper, divided into four main parts: Data acquisition with three methods; pre-processing; dynamic modeling and finally validation between the three abovementioned measures.

### 2.1 Field Site and Experiments

All experiments were conducted at the ETH Zürich research station for Plant Sciences Lindau-Eschikon, Switzerland (47.449 N, 8.682E, 520 m a.s.l.) within or right next to the Field Phenotyping Platform (FIP) (Kirchgessner et al. 2017). The experimental field is a gleyic cambisol soil with 21% clay, 21% silt and 3.5% organic matter (Kirchgessner et al. 2017). Data originating from four different experiments were used (Table 1). RGB and SPC measurement were acquired for three experiments (2018, 2021 and 2022 all FIP main) and visual scoring for two experiments (2018 and 2021) respectively.

**Table 1.**
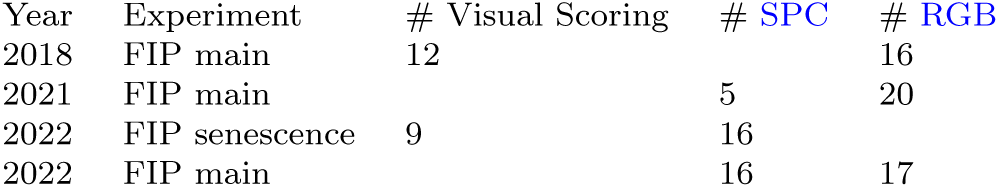
Overview of experiments. The *FIP main* in 2018 is described in more detail in Anderegg, Yu, et al. 2020 and the *FIP senescence* experiment in 2021 in Anderegg, Zenkl, et al. 2023

| Year | Experiment | # Visual Scoring | # SPC | # RGB |
| --- | --- | --- | --- | --- |
| 2018 | FIP main | 12 |  | 16 |
| 2021 | FIP main |  | 5 | 20 |
| 2022 | FIP senescence | 9 | 16 |  |
| 2022 | FIP main |  | 16 | 17 |

### 2.2 Visual Scoring

Reference assessments of senescence were obtained through visual inspection of canopies at regular intervals, starting before the onset of senescence in the earliest-maturing genotype and continuing until the latest-maturing genotype reached maturity. Canopies were inspected at a view angle of approximately 45°, lower leaf layers were manually exposed where necessary to enable inspection. All estimates were made for a central region of about 0.5 m x 0.5 m in each plot. In 2019, canopy senescence was assessed by estimating the fraction of green, chlorotic, and necrotic leaf and stem area, with chlorotic areas interpreted as representing an intermediate stage of senescence. The resulting average impression of the advancement of senescence was expressed as an integer value on a scale ranging from 0 (representing complete canopy senescence) to 10 (representing fully stay-green canopies). Ear senescence was not considered. In 2022, the focus of the visual scorings was put on the remaining green leaf area, whereas stem and ear senescence was not considered, and no distinction was made between chlorotic and necrotic tissue, with both considered as “senescent”. All visual scorings were made by a single person to avoid scorer-bias. Data was recorded using the field book app (Rife et al. 2014).

### 2.3 Sensors

For RGB image acquisition, a camera with a full frame sensor of 6000×4000 pixel (Sony *α* 9 Model ILCE-9, Sony Corporation, Tokyo, Japan) equipped with a prime lens with a focal length of 55 mm and a maximum aperture of f/1,8 (Sonnar T *∗* FE 55 mm F1,8 ZA, Sony Corporation, Tokyo, Japan) was used as described in Roth, Camenzind, et al. 2020. For the SPC image acquisition a 10 band multispectral sensor (Micasense Red Edge MX dual cam system, MicaSense, Seattle, United States) was used (see Table A1). Both sensors were mounted to a Matrice 600 Pro drone (SZ DJI Technology Co. Ltd., Shenzhen, China) and attached to a Ronin-MX gimbal (SZ DJI Technology Co. Ltd., Shenzhen, China) in order to minimize measurements with off-nadir position. The used multispectral and RGB cost approximately 12’0000 and 1’500 USD, respectively. Cameras with comparable specifications and especially resolution can be found for less than 1000 USD. This means that the RGB camera is currently between eight and eleven times cheaper than the SPC sensor.

### 2.4 UAV Measurements

The RGB-UAV flights were performed at 28 m altitude with front overlap ratio of 90 % and a side overlap ratio of 80 %. This resulted in a ground sampling distance (GSD) of 3 mm (Roth, Hund, et al. 2018). The experimental field was equipped with ground control point (GCP)s arranged cross-wise with a spacing or 12 m *×* 18 m, according to Roth, Camenzind, et al. 2020.

The SPC campaigns were conducted according to Perich et al. 2020 with a flight height adjusted to 100 m and the above described SPC sensor. The resulting GSD for the SPC images was *∼* 5 cm. We aimed for a minimum of two measurements per week. However, measurements with the multispectral sensor require specific measurement conditions (very stable light conditions in particular). Therefore, maintaining a rigid measurement schedule was not always possible.

### 2.5 Image pre-processing

RGB and SPC images were pre-processed in a similar pipeline using Agisoft PhotoScan Professional 1.4.2 and Agisoft Metashape 1.5.2 (Agisoft LLC, St. Petersburg, Russia) and OpenCV (Bradski 2000). Images from a given measurement flight were first aligned, and camera position estimations referenced using GCPs placed in the field. The world-coordinate positions of the GCPs were measured with a Trimble GPS RTK (R10, Trimble Ltd., Sunnyvale, U.S.A.) with a maximal error of 2 cm. From a generated dense point cloud, a digital elevation map was extracted and orthophotos were projected onto it. This orthophoto was used only to define the region-of-interest (ROI) of plots, but not for data extraction. To avoid fusing pixels from different shots and height-related artifacts, these ROIs were back-projected to the original images (according to Roth, Aasen, et al. 2018; Roth, Hund, et al. 2018). Plot-values were then extracted from the most nadir-view image per plot.

#### 2.5.1 Multispectral calibration

In order to reduce the effect of varying environmental conditions such as changes in lighting across individual flights, we conducted a calibration. Calibration of the SPC images was done after the clipping of individual plots from orthophotos. At the beginning of each flight, five control panels with a known reflectance (0 %, 15 %, 25 %, 75 % and 100 % black) were measured. Measured panel reflectances were then used to fit a linear model according to the empirical line method (Baugh et al. 2008), and predict the calibrated multispectral values. Calibrated multispectral reflectances were used for all further analyses.

### 2.6 Reflectance Index calculation

For the calculation of reflectance indices (see Table A2 and A3) the reflectance values of a central region of interest within each plot were considered. The regions of interest were defined by maintaining a margin of approximately one sowing row (12.5 cm) on each side of the plot to reduce border effects. Within the region of interest the index was calculated for each pixel and the median of the resulting index values was exported for further analyses. 25 and 16 indices were extracted from the SPC and the RGB imagery, respectively.

### 2.7 Dynamic modelling

Various approaches for modelling the dynamics of senescence based on repeated visual scoring, measurements of green leaf area, or spectral indices have been proposed, and different sets of parameters are often extracted from model fits (Chapman et al. 2021). The choice of model and parameters depends on the level of assessment and on the frequency with which these assessments are made. At the canopy level, senescence typically shows a sigmoidal progression, which can be well represented by two-parameter or four-parameter logistic-type models. Alternatively, splines may provide more flexibility to represent atypical senescence curves as well, or may be used to model spectral indices that deviate in their dynamics from visual observations. However, using more flexible approaches increases the risk of overfitting to errors associated with single measurements or scorings when compared to more rigid fully parametric approaches. Here, we evaluated both approaches. Four-parameter Gompertz models were fitted as described earlier (Anderegg, Yu, et al. 2020):

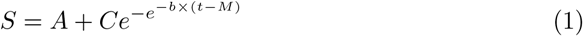

where *S* represents the trait value; *A* and *C* are the lower and upper asymptotes, respectively; *b* is the rate of change at time *M*; and *M* is the time point when the rate is at its maximum (Gooding et al. 2000). A corresponding two-parameter model with asymptotes constrained to 0 and 10 was also fitted. Monotonically decreasing P-splines were fitted using the R-package scam (Pya et al. 2014), with the number of spline knots set to three quarters of the number of observations, as done for similar data by Roth, Rodríguez-Álvarez, et al. 2021. Given that our main interest is in extracting the dynamics of senescence rather than absolute values at a specific point in time, all scorings and measurements were scaled at the plot level to range from 0 to 10, representing the minimum and maximum value recorded for the assessment period, respectively. To simplify modelling and model interpretation, the scale for indices with increasing values during senescence was inverted. The different models were applied to the visual scoring as well as to all indices. The correlation between visual scoring and the high throughput field phenotyping (HTFP) method was then calculated using the equivalent model for each (see Figure 1).

### 2.8 Extracting key time points

From the fitted models, a value was predicted for each plot on each day in the assessment period. The predicted values were used to extract the parameters *t*20, *t*50 and *t*80 as the time points when modelled scaled scorings or index values decreased to 8, 5, and 2, i.e., when values decreased to 80 percent, 50 percent, and 20 percent of the maximum observed value within the assessment period. Additionally, the *Integral* under the fitted curves was extracted. For fully parametric models, we also extracted the *M* parameter (see above) directly from the model fit. Finally, the onset (*onsen*) and endpoint (*endsen*) of senescence were extracted from parametric models as the time point when the second derivative was at its minimum and maximum, respectively (Figure 2). Methods for dynamic modeling and key parameter extraction can be found in the public open git repository.

**Fig. 2.**
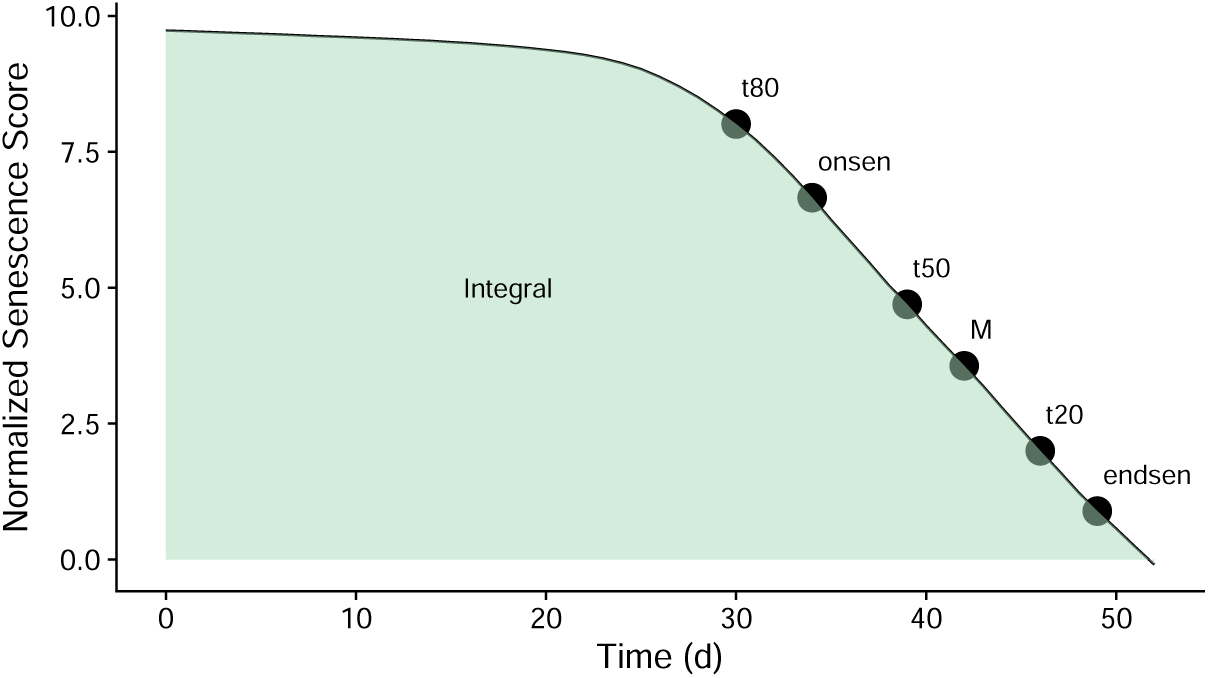
The figure shows the seven extracted parameters laying on a index trajectory (x-axis represents time, y the normalized senescence score value). Six of the parameters are time points (*t*80,*onsen*,*t*50,*M*, *t*20 and *endsen*) and one describes the area below the index curve in green (*Integral*).

### 2.9 Best index selection

The best index-model combination was selected based on the dynamic modeling and determination of the key time points. For each parameter-index-model combination, the correlation score between the visual score and the HTFP was calculated. First, the median correlation of all selected parameters per index was calculated, without reference to the underlying models. In addition, the range of correlation values was taken into account by selecting the best index. That is, a slightly lower correlation value was preferred if the range between the minimum and maximum value was smaller. A smaller range represents a more robust index and was therefore considered superior in this study. In a second step, the best model was selected according to the same metrics as before, but calculated per model within the previously selected best index per sensor. This method allows to determine the index-model combination that represents the best correlation and therefore the relative behavior of the genotypes, but does not take into account the absolute deviation or bias between two compared methods. For the overall comparison of trait extraction and correlation, the best index-model combination was also selected for each selected parameter.

### 2.10 Sensor Comparison

The comparison between RGB and SPC measurements was based on the parameters *t*20, *t*50 and *t*80. These parameters were chosen because they are less affected by a late start of measurements or too early ending. For each of the parameters, the best index-model combination of each sensor was taken and the Pearson correlation as well as the root mean squared error (RMSE) was calculated.

### 2.11 Temporal measurement density effects

To evaluate the effect of measurement frequency on the estimation of senescence dynamics parameters, we generated subsets of the full data set that contained a decreasing number of measurement time points (n). Nine independent subsets were sampled per n, with n=5, 6, …, 16. This procedure was repeated twice: Once enforcing the first and last measurement time point to be contained in the subset, and once relaxing this constraint. Each resulting subset was then subject to dynamic modelling and parameter extraction as described above. The analysis was performed for the best index-model combination identified previously.

## 3 Results

We used a dataset containing four different experiments that were conducted over three years to evaluate the potential of UAV-based spectral and RGB imaging to capture the senescence process in wheat. This comprehensive dataset represents more than 2000 individual plots and a large genotypic diversity, providing a vast basis for our analyses. In order to compare the two sensors, we included data from two experiments which were measured with both sensors. This facilitated the comparison of the sensors, but also allowed us to explore potential year-to-year variations in the data, providing valuable insight into robustness over the years of features derived from each of the sensors.

First, the best index was selected for each sensor. Then, each model was applied to the visual scoring and the indices, and the resulting parameters were correlated. For visual scoring, the type of model applied had a negligible effect on the estimation of the senescence dynamics parameters (all pairwise correlations *>* 0.97; Figure A8 and Table A6). For SPC *NDV Ired*650 and for RGB *ExGR* emerged as the best indices (Figure 3). For the SPC indices, *NDV Ired*668 had a slightly higher median correlation but a much wider range of correlation values across all parameters of interest than *NDV Ired*650, so the latter was preferred. *ExGR* was clearly the best RGB index in both aspects. Overall, the *cgom* (C-Gompertz) model provided the best approximation of (Table A4). Therefore, this model was selected for all further analyses. The *cgom* model appeared to be more meaningful than the *fgom* model in following the actual senescence dynamics. While the *fgom* has more degrees of freedom, it can follow measurement errors more directly (Figure 4; right side), the same is true for the *pspl* and *lin*. This is a problem that only occurs with unclear index trajectories or larger uncertainties (Figure 4; good versus bad plots), otherwise, the different models do not differ much (see also Figures A1, A2, A3 and A4).

**Fig. 3.**
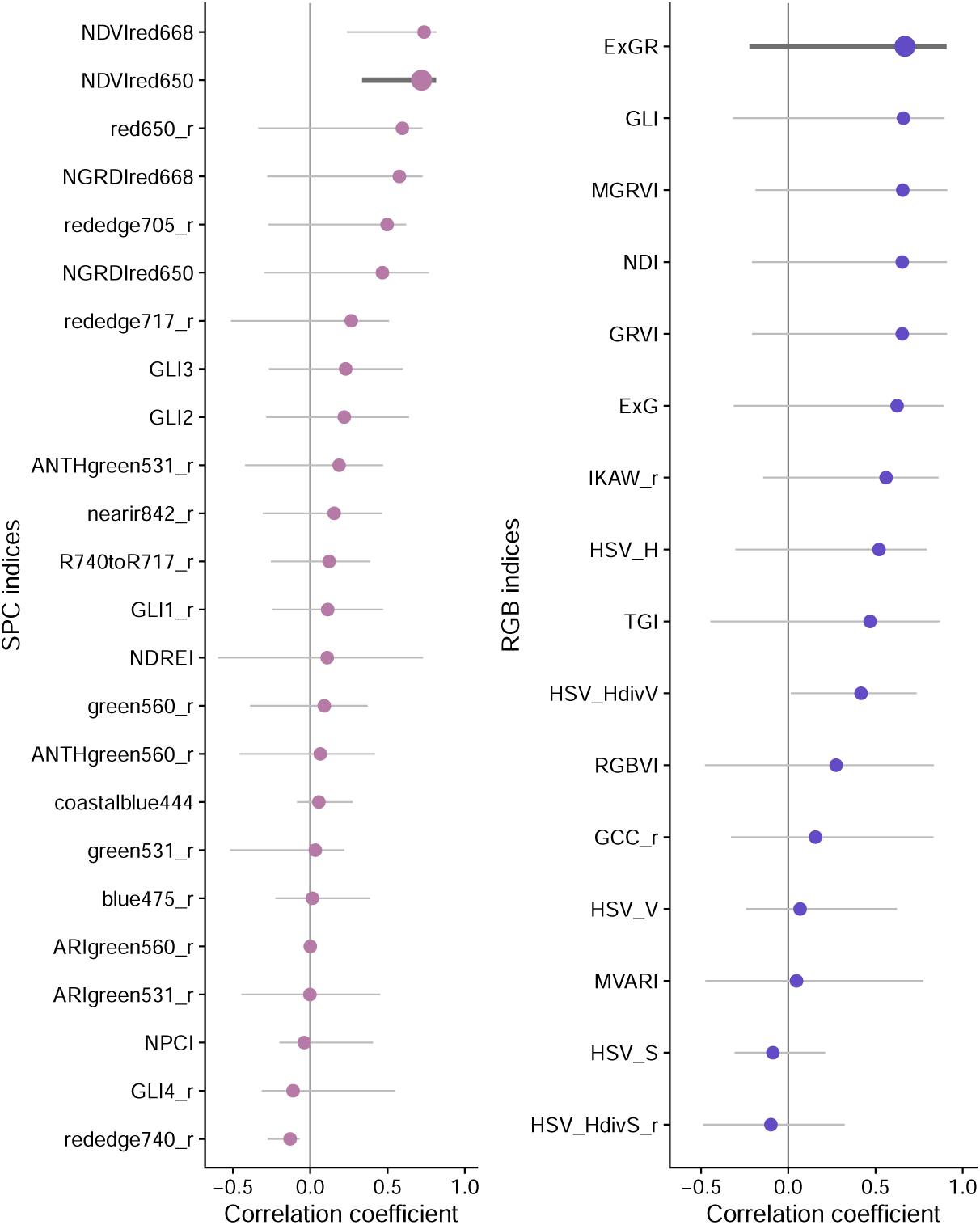
Selection of the best index for SPC (left) and RGB (right) was done according to the median (points) and the range (minimal and maximal - whiskers) correlation values over all used parameters. Thicker line represents the best index for SPC the *NDV Ired*650 and for RGB the *ExGR* index have been chosen.

**Fig. 4.**
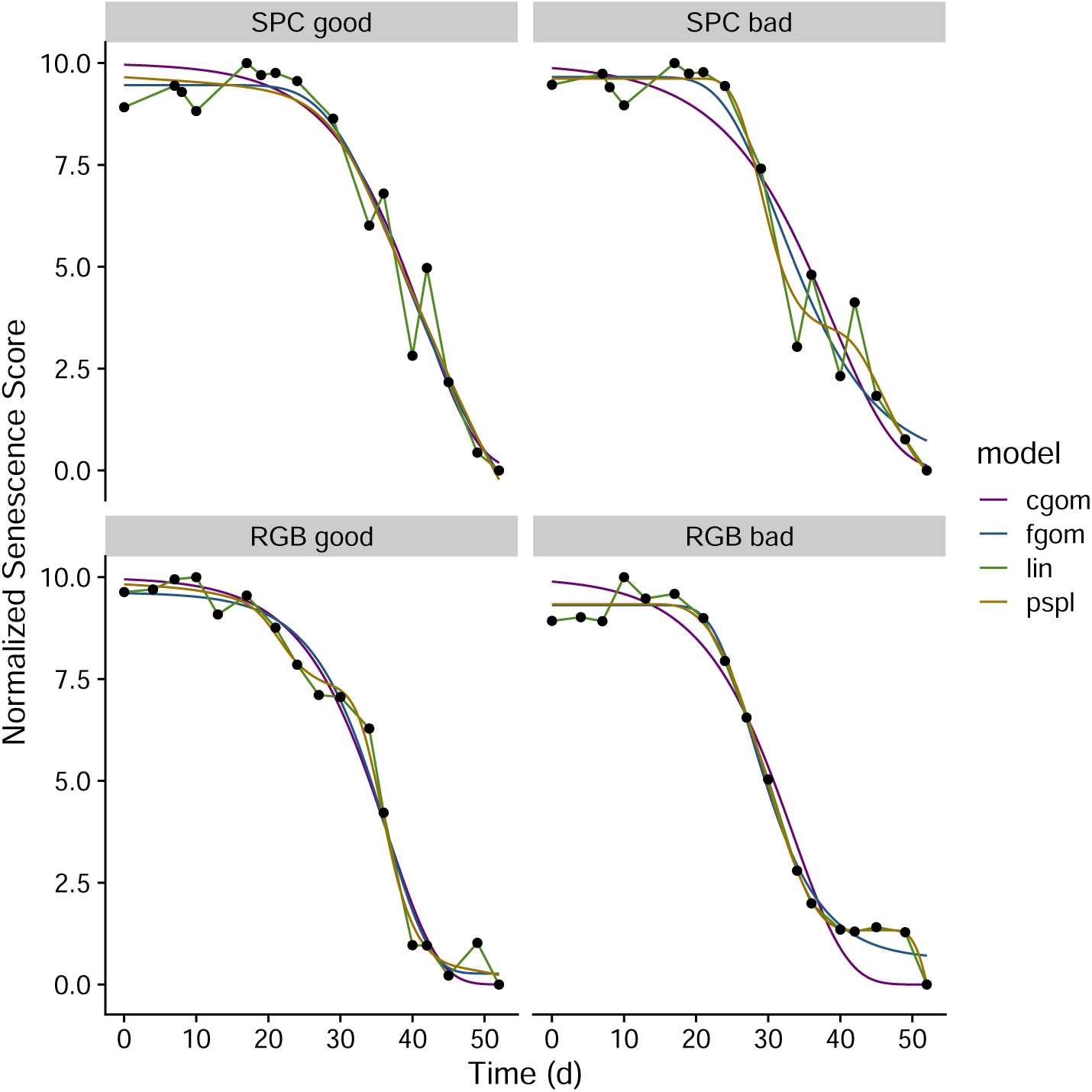
For two experimental plots the different model fits are shown (lines), in the upper row for the selected SPC index (*NDV Ired*650) in the lower row for the selected RGB index (*ExGR*). The points represent the scaled measurement index values.

Whereas several parameters correlated well between SPC indices and visual scoring (*t*80, *t*50, *onsen*, *Integral* and *M*), weaker correlations were also observed for some parameters, i.e. *t*20 and *endsen* (Figure 5) by using the *NDV Ired*650-index with the *cgom* model. For the parameter *Integral* a decent correlation but a substantial bias was observed. When considering the best index-model combination for each parameter individually, correlation coefficients ranged from from 0.73 (*t*80) to 0.82 (*t*80) (Figure A5). For five of the seven parameters the *NDV Ired*650 or *NDV Ired*668 were the best indices. However, just in two occasions (*Integral* and *onsen*) in combination with the *cgom* model.

**Fig. 5.**
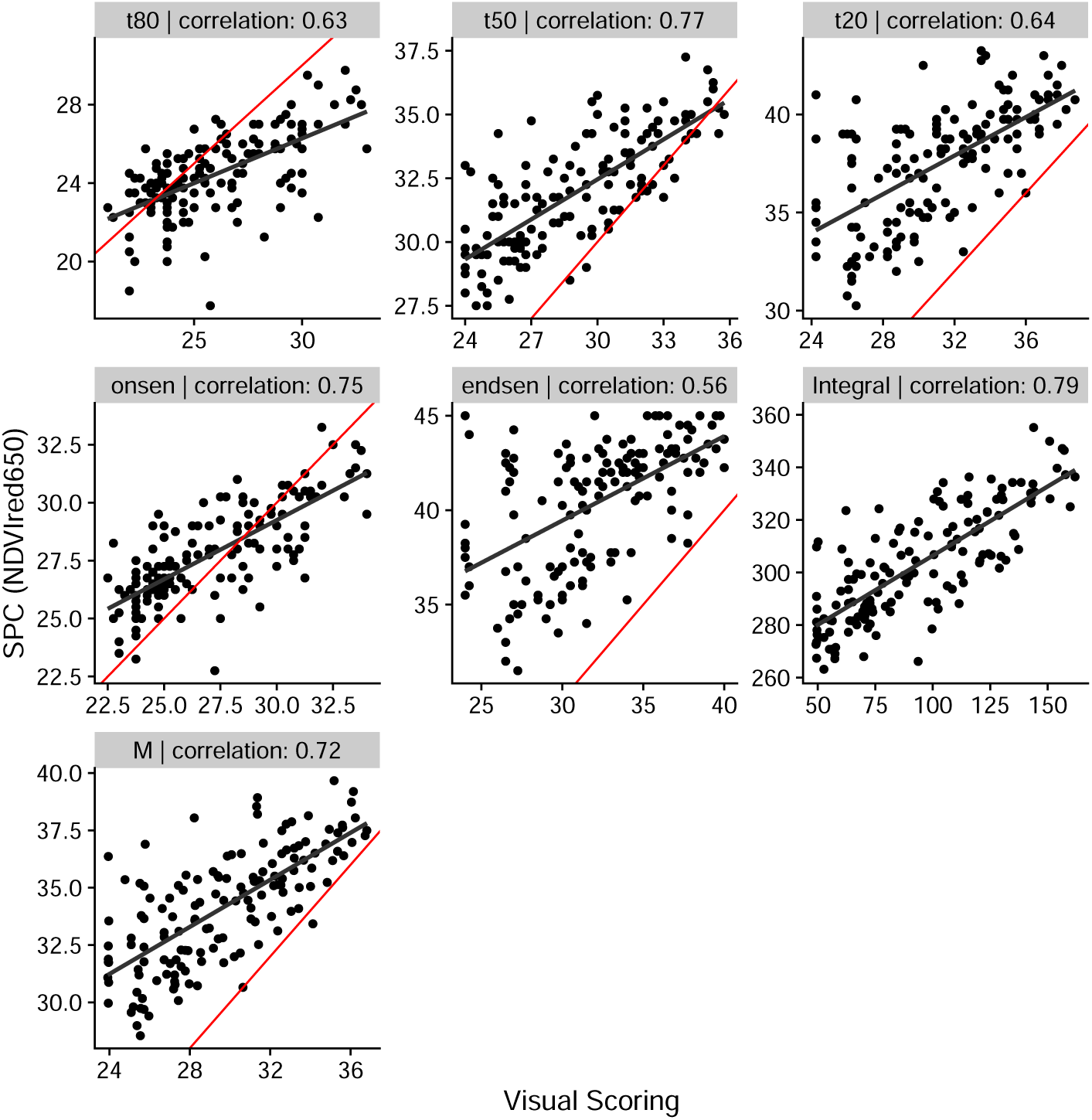
Correlation between visual scoring (x-axis) and the best SPC index (*NDV Ired*650 with the *cgom* model) measured by High Throughput Field Phenotyping (HTFP). The red line represents the 1:1 line, while the black line shows the correlation fit. Each panel shows one parameter (indicated in the header) and the corresponding Pearson correlation value.

For the RGB indices, the *ExGR* with the *cgom* model showed overall the best performance (Figure 3 and Table A4). Again, most of the parameters show a strong correlation between RGB indices and visual scoring (*t*50, *t*20, *endsen*, *Integral*, and *M*), while *t*80 and *onsen* do not show a strong or any correlation (Figure 6). By selecting for each parameter the best combination, we found correlation coefficients ranging from 0.32 (*t*80) to 0.91 (*t*20) (Figure A6). The *ExGR* index was found twice to be the best index per parameter as well (*Integral* and *M*), however just once in combination with the *cgom* model.

**Fig. 6.**
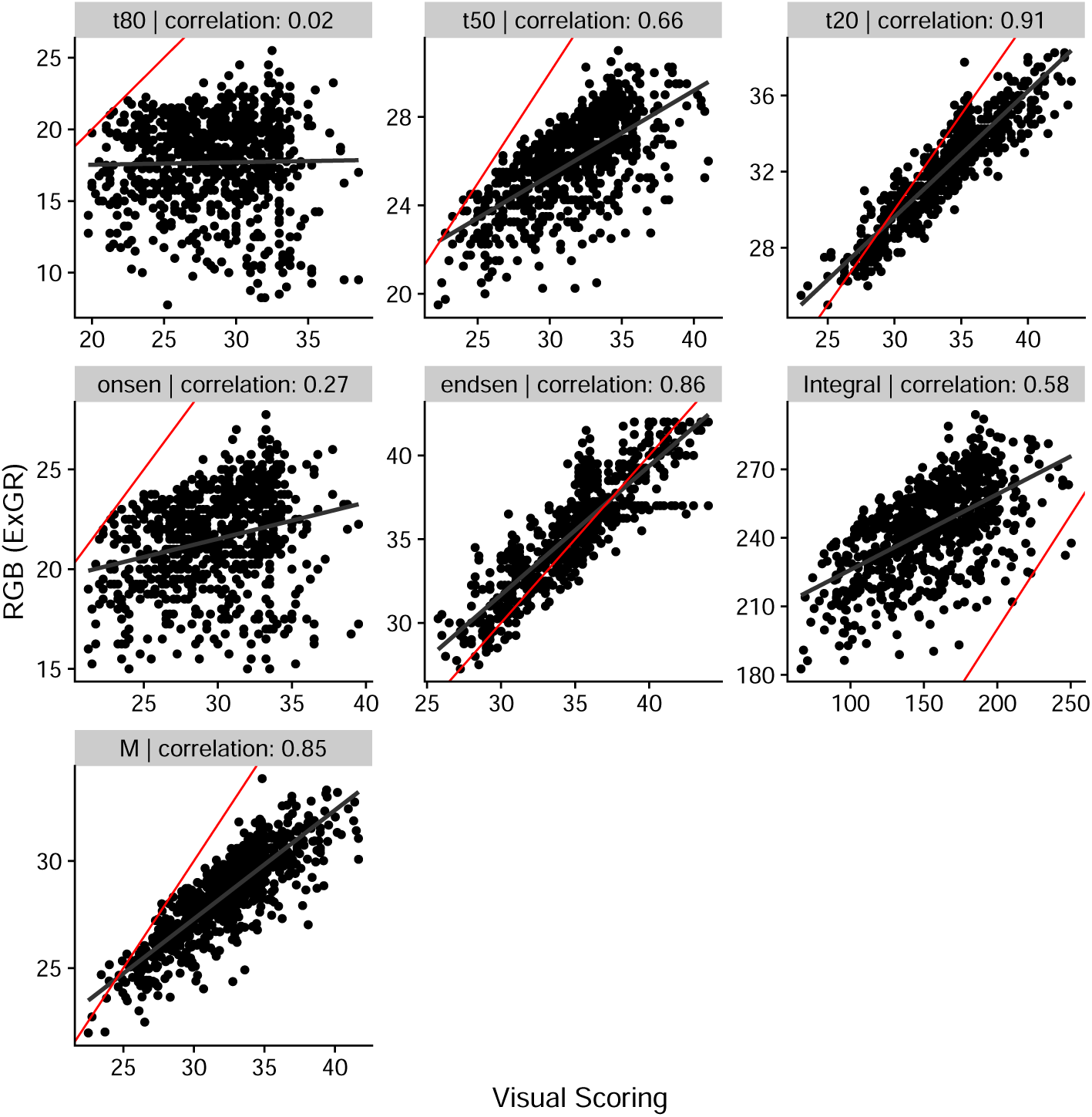
Correlation between the visual scoring (x-axis) and the the best RGB index (*ExGR* with the *cgom* model) measured by high throughput field phenotyping (HTFP) (y-axis). The red line represent the 1:1 line, whereas the black line shows the correlation fit. Each panel shows one parameter (indicated in the header) and the corresponding Pearson correlation value.

Dynamics parameters derived from the best RGB and SPC indices correlated over two years of experiment (Table 1, FIP main 2021 and 2022). Years were analyzed separately, allowing to check for effects deriving from the year. Correlation coefficients all showed values between 0.55 and 0.72 in 2022. In 2021, correlation values ranged from −0.02 for *t*80 up to 0.4 for *t*50 (Figure 7). Zero values can occur if a model converges but predicted values never cross the pre-defined thresholds required for parameter extraction. The resulting correlation per year with all the parameters as well as RMSE values are shown in Table A5.

**Fig. 7.**
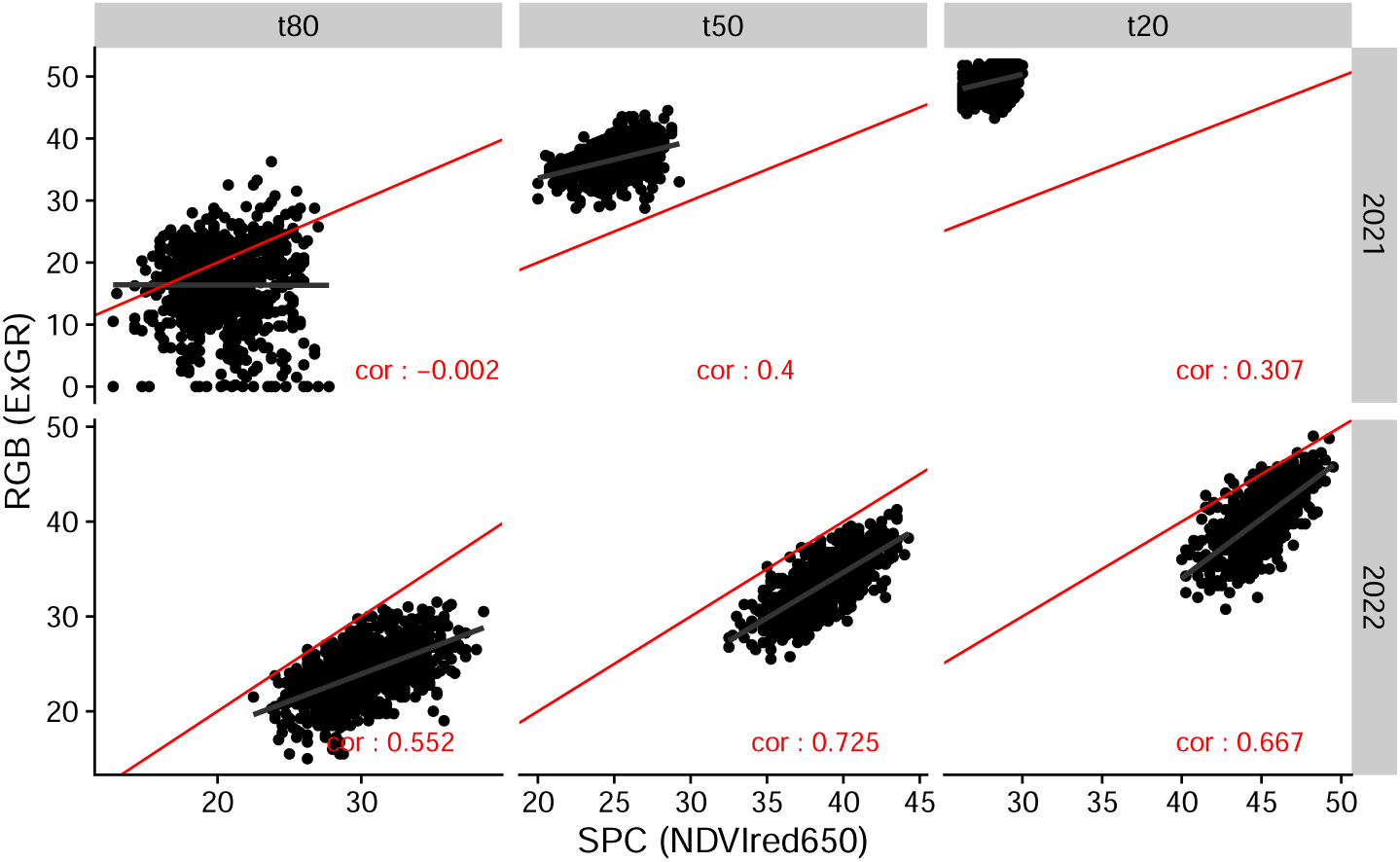
Correlation between the best RGB index-model combination (*ExGR* - *cgom*; x-axis) and the the best SPC index-model combination (*NDV Ired*650 - *cgom*; y-axis). The red line represent the 1:1 line, the black line shows the correlation fit. The upper row shows the data measured in the year 2021 the lower the data of 2022. The columns represent the different parameters (*t*20, *t*50 and *t*80), in each panel the corresponding Pearson correlation value is written in red (see Table A5)

Taken together, the extracted parameters explain and define the dynamics of the senescence process. However, the overall dynamics of the process can also be visualized as continuous over time. Figure 8 shows the median of the best index across two years (2021 and 2022) of SPC and RGB over time. In 2021, the SPC measurements started 19 days after the RGB campaign. The parameter *t*80 derived from the RGB measurement in 2021 is in the phase where no SPC measurement took place. In 2022, the measurement period with both sensors was equivalent in duration. Both trajectories show a very similar trend, a relatively long and pronounced initial phase with values around 10. Senescence then starts slowly, visible by the decreasing value, followed by a linear phase. This phase either ends abruptly due to the harvest, or the slope is again slightly reduced towards the harvest time or towards the end of the measurement phase.

**Fig. 8.**
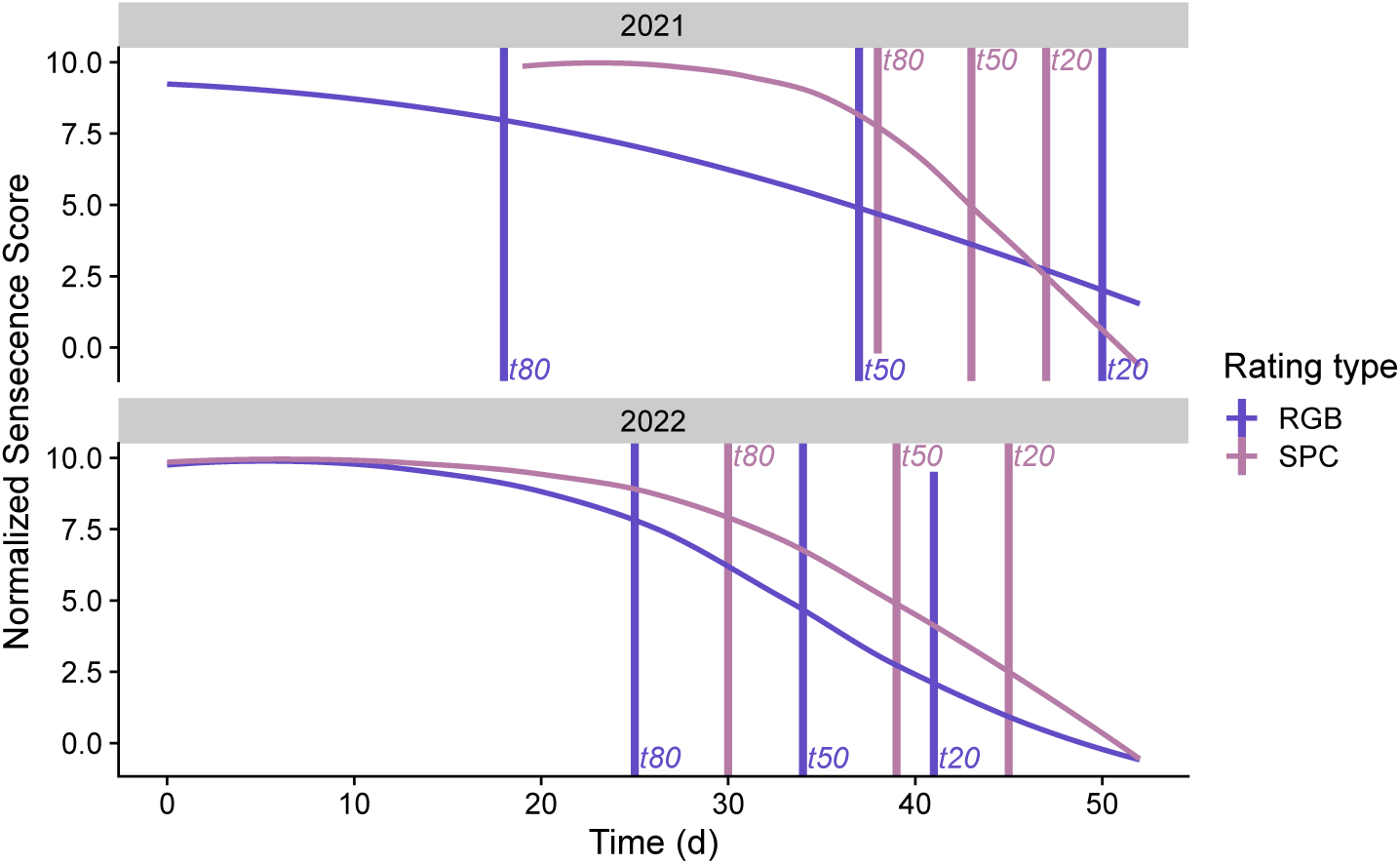
senescence dynamics as measured via RGB and SPC (y-axis median of the standardised senescence score of all measured plots) over time (y-axis), starting from the first day of measurement (0) until the last day (53), in two different years (upper panel: 2021, lower panel: 2022). Vertical lines represent the median of parameter *t*80, *t*50 and *t*20 per year measured by the different sensors.

The accuracy of extracting the parameter shown is highly dependent on the data at hand. Subsampling the measurement series generally resulted in decreased correlation coefficients and higher variance in estimated parameters (Figure 9). The overall tendency in both sampling strategies shows lower correlation coefficients and especially a wider spread of the values when reducing the measurement times. Parameters at the very beginning or end of the senescence phase (e.g. *t*80 and *t*20) show less reduction in accuracy and variance than others (e.g. *Integral*). The performance with fixed start and stop values seems to be overall better and more stable (lower variance) than with unconstrained subsampling. In particular, the SPC sensor shows a large increase in variance and a reduction in correlation coefficient overall with a lower number of measurements.

**Fig. 9.**
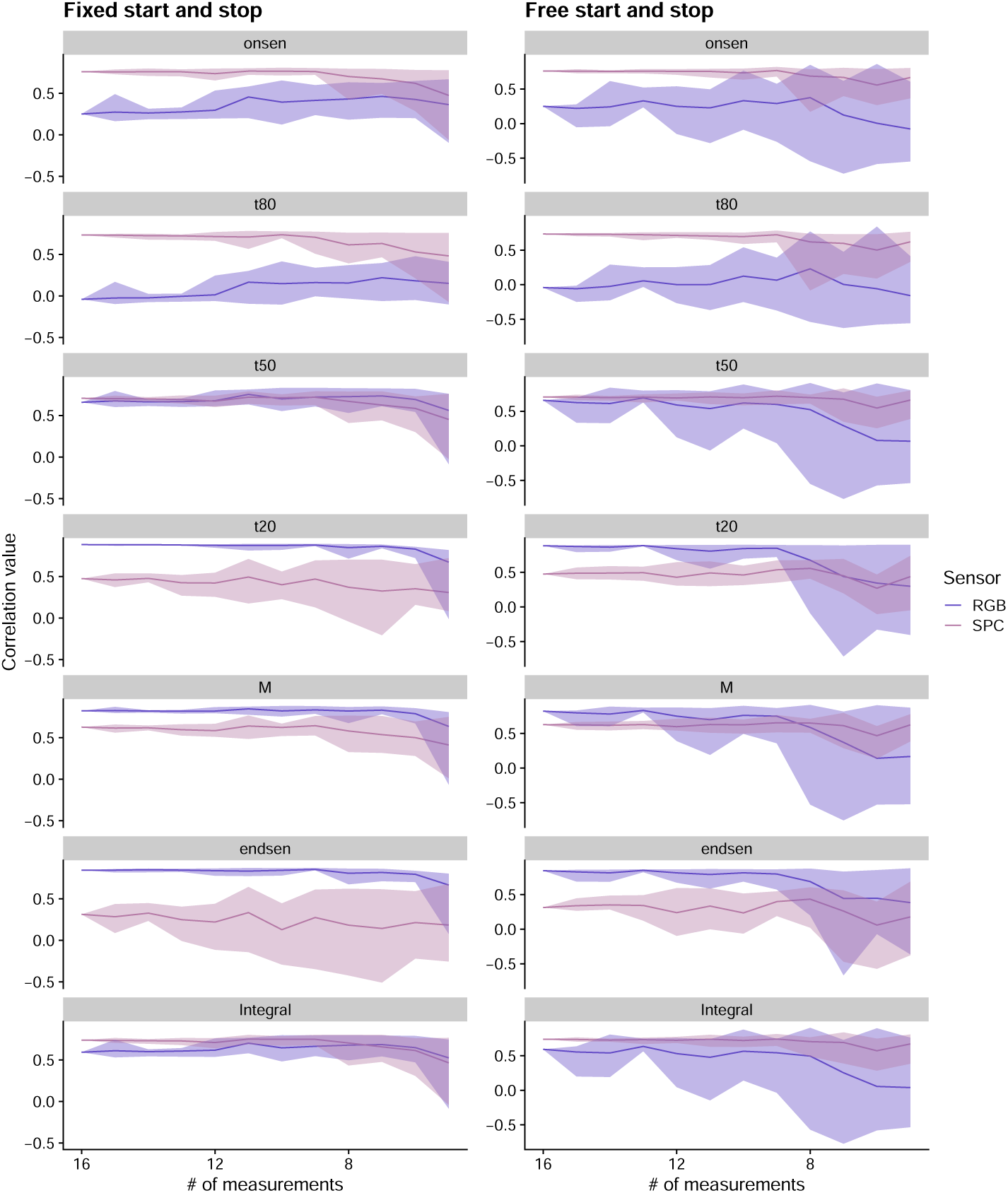
Correlation coefficients (y-axis) over the decreasing number of measurement time points (x-axis). Each row shows one of the parameters described above. The row represents the different sub-sampling approaches, right with maintaining start and stop of the measurement phase, left without maintaining start and stop but freely reducing the sample size. The different colors indicate the two sensors (RGB and SPC), the line represents the median, the shades the minimal and maximal values.

## 4 Discussion

This study transfers a methodology for extracting key parameters of the senescence phase from ground-based to UAV-based reflectance measurements, providing the required high throughput Anderegg, Yu, et al. 2020. Extensive data sets from multiple years allowed us to thoroughly validate a SPC and a RGB sensor as tools to track visually observed canopy senescence. For both sensors we identified a robust reflectance index that allows to track this process, namely for the SPC the *NDV Ired*650 and for the RGB sensor the *ExGR* index in combination with a dynamic model (*cgom*). By validating these sensors against visual scoring, the state of the art in plant breeding, as well as against each other, we could show a strong correlation regarding several parameters (Figure 5 and 6). We propose a robust method to extract key parameters of the senescence phase in a high throughput manner. Furthermore, our results indicate that the accuracy of assessments depends primarily on the temporal resolution of the measurements, rather than on the availability of narrow-band, specific spectral indices. This suggests to invest more in a temporal higher resolved data set than a spectral higher resolving sensor.

The described method allows the up-scaling of previously developed approaches, such as Lopes et al. 2012; John T. Christopher, Veyradier, et al. 2014; Anderegg, Yu, et al. 2020, allowing such methods to be implemented in actual breeding programs due to a feasible amount of work required to collect the necessary datasets. Furthermore, our results show that both evaluated sensors can deliver similar results (Figure 7; Table A5) and provide a representation of the senescence process that is similarly accurate to that obtained by using more sophisticated sensors, such as spectroradiometers (Anderegg, Yu, et al. 2020). This indicates that cheaper RGB sensors can be used similarly, in line with results of Cao et al. 2021. Also, such RGB sensors are widely used for other phenotyping purposes (Ge et al. 2016; Jin et al. 2017; Roth, Camenzind, et al. 2020; Anderegg, Tschurr, et al. 2023). Compared to SPC sensors, RGB sensors are more robust regarding different illumination conditions, and provide often higher resolution but fewer measured bands.

However, this also means that measurements can be made under less optimal conditions, which can facilitate more continuous monitoring of experimental plots. Our results confirm that a high measurement frequency and a sufficient duration of measurement campaigns are more important to collect valuable information than the chosen sensor, and particularly, its spectral resolution. Figure 8 shows a very late start of SPC measurements in 2021, which is also reflected in a low correlation (correlation coefficient: 0.2; see also Table A5) between RGB and SPC for *t*80, a parameter defining the early phase of senescence. When the measurement period was identical (as in 2022) both sensors are equal in accuracy, which is also reflected in high correlation coefficients (Table A5). When considering the best index per parameter for each sensor, higher correlation coefficients were observed (Figure A5 and A6). The indices show a slightly different pattern over time (Figure A7). This may explain why the parameters can vary between the indices, and thus the correlation between the visual scoring and the HTFP derived measurement can be increased by selecting an index-model combination separately for each parameter. However, as can be seen in Figure 4, there are also differences between the different models used. The differences are small if the period was measured in high temporal resolution. In this case, the measurement error is also small (see also Figures A1, A2, A3 and A4). If this is not the case, some models or indices follow a pattern that is not physiologically meaningful. Therefore, if one is interested in observing the whole dynamic of a period, and may compare it between different environments, we suggest using one index with one model, to be chosen as described in this study. This allows a robust tracking of the phase of interest. Conversely, if the interest lies in extracting a single key time point, the precision can be maximized by selecting a specific index for each index. In our study, selecting the best index-model combination per parameter increased the correlation between 0 and 0.48 for RGB and 0.01 to 0.19 for SPC (see Figure 6 versus A6, respectively A5 versus A5).

As shown in Figure 3, the different models also have an effect on the correlation score. While there is often little difference between the two types of Gompertz models, there is an overall tendency for both P-Splines and the linear model to perform worse. This indicates that fully parametric approaches are better suited to model canopy senescence dynamics.

The temporal density of the measurements was shown to have a greater effect on the results and on the variance of the extracted parameters than the sensor (Figure 9). However, some parameters can be extracted with comparable precision and variance even with a reduced number of measurement points (e.g. *t*80, *t*50, and *t*20) if the start and end of the period are fixed to a physiologically meaningful point in time. If this is not the case (Figure 9 right side), the correlation coefficients show a higher variance with reduced measurement times. This trend can be observed for both sensors. Therefore, for a robust and meaningful parameter extraction and a better understanding of the observed period, continuous measurements over time as well as a meaningful timing regarding the start and end of the measurement campaign are key. There are proposals to interpolate the measured features using an environmentally meaningful component (e.g. temperature) to fill measurement gaps as meaningfully as possible (Graf et al. 2023). However, this requires a huge amount of data to train on a genotype-specific level, which is unlikely to be available if the measurements were not made in a dense temporal resolution in the first place. With more bands available in the SPC sensor, more potential combinations for indices are possible. This may be of interest not only to follow the dynamics of senescence, but also to study the contribution of biotic and abiotic stress to the overall dynamics of green leaf area (Anderegg, Hund, et al. 2019).

The ability to measure at high temporal resolution allows for a deeper understanding of senescence dynamics. These dynamics can then be used to identify potential stressors, similar to what was done in Tschurr, Kirchgessner, et al. 2023 for dynamic canopy cover measurements. And in a next step, implemented in crop models to further improve prediction accuracy by accounting for growth stress according to measurable traits (Diepen et al. 1989; Wang et al. 1998; Jamieson et al. 1998; Brown et al. 2018). This approach could also be used to assess and improve the performance and phenological adaptations of crop mixtures in a high-throughput manner (Tschurr, Oppliger, et al. 2023).

The RGB sensor already has a higher GSD than the SPC sensor, but for both the resolution was not sufficient to distinguish between plant and soil pixels in the images during the senescence phase. During earlier growth phases, the resolution of the RGB sensor was high enough to allow this (Roth, Camenzind, et al. 2020). In the senescence phase this requires very high resolution, but would allow to determine the senescent leaf area Anderegg, Zenkl, et al. 2023. This could ultimately allow to determine the actual state of senescence, by defining the ratio between green and already dead plant tissue. These measurements could then also be modelled and evaluated with the suggested method in this study, treated as one of the shown indices. Furthermore, even organ-level senescence dynamics could be extracted when the resolution is sufficient. This could allow to disentangle flag leaf senescence, whole plant senescence, grain maturation and their corresponding genotypic patterns (Chapman et al. 2021). Lower UAV flight height as well as an adapted flight strategy (i.e. Xiao et al. 2023) with a high resolution RGB camera could allow this. However, the resolution of SPC sensors is typically lower, again suggesting further investigation and development of methods using RGB sensors.

## 5 Conclusion

This study highlights the promising potential of up-scaling and improving the efficiency of senescence measurements by the use of UAV. Our results show comparable results to those obtained by previous state-of-the-art methods. In addition, we propose a strategy that prioritizes increasing measurement frequency over time, as opposed to investing in more expensive sensors equipped with a high number of spectral bands. Our investigation showed similar results when using a standard RGB sensor as opposed to a 10-band SPC sensor, consistent with previous research using hyperspectral sensors. Particularly for the senescence phase of winter wheat, a phase characterized by substantial genotype by environment (G*×*E) interactions, achieving high temporal resolution and measurement throughput is crucial to advance our understanding and selection of well-adapted genotypes.

## Acknowledgments

We thank Hansueli Zellweger and Simon Corrado for managing our field experiments over the years, and members of the Crop Sciences group for valuable help in data acquisition and fruitful discussions. Furthermore we want to thank valuable help from Price Akiina in data collection as well as all of the Crop Science Group of ETH Zurich.

## Author Contributions

FT: Conceptualization, methodology, software, formal analysis, visualisation, data acquisition, writing (original draft)

LR: Data acquisition, supervision, writing(review and editing)

NS: Methodology, software, data acquisition, writing (review and editing)

OZ: Data acquisition, writing (review and editing)

AW: Funding acquisition, writing (review and editing)

JA: Conceptualization, supervision, methodology, software, formal analysis, data acquisition, writing (original draft)

## Funding

FT and LR were founded by the Swiss National Science Foundation Grant No. 200756.

## Conflicts of Interest

The authors declare that they have no competing interests.

## Data Availability

Code and example data available at: dynamic senescence modeling.git

## Appendix A Supplementary Information

### A.1 Supplementary Tables

**Table A1.** Bandwiths of the Micasense Dual Camera system.

| Band Name | Center Wavelength (nm) | Bandwidth (nm) |
| --- | --- | --- |
| blue475 | 475 | 32 |
| green560 | 560 | 27 |
| red668 | 668 | 14 |
| rededge717 | 717 | 12 |
| nearir842 | 842 | 57 |
| coastalblue444 | 444 | 28 |
| green531 | 531 | 14 |
| red650 | 650 | 16 |
| rededge705 | 705 | 10 |
| rededge740 | 740 | 18 |

**Table A2.** The used RGB-indices are shown in the table below, whereas r refers to the red-color band, g to the green and b to the blue one. Furthermore the Hue, Saturation, Value color space (HSV) was used whereas the h stands for the hue value, s the saturation and v for the value. In the TGI following *λ*-values have been used: *λ_r_*= 670; *λ_b_* = 480; *λ_g_* = 550.

| Index | Formula | Reference |
| --- | --- | --- |
| TGI | $-0.5 \times ((\lambda_r - \lambda_b) \times (r - g) - (\lambda_r - \lambda_g) \times (r - b))$ | used inAnderegg, Tschurr, et al. 2023 |
| ExG | $2 \times g - (r + b)$ | used inAnderegg, Tschurr, et al. 2023 |
| NDI | $128 \times (\frac{g-r}{g+r} + 1)$ | used in Anderegg, Tschurr, et al. 2023 |
| ExGR | $(2 \times g - (r + b)) - (1.3 \times r - g)$ | used as in Anderegg, Tschurr, et al. 2023 |
| GCC | $\frac{g}{b} + g + r$ | used in Yue et al. 2017 |
| GRVI | $\frac{g-r}{g+r}$ | used in Jing et al. 2017 |
| IKAW | $\frac{r-b}{r+b}$ | used in Maimaitijiang et al. 2019 |
| MGRVI | $\frac{g^2-r^2}{g^2+r^2}$ | used in Cen et al. 2019 |
| MVARI | $\frac{g-b}{g+r-b}$ | used in Cen et al. 2019 |
| RGBVI | $\frac{g^2-b \times r}{g^2+b \times r}$ | used in Possoch et al. 2016 |
| HSV_H | $h$ | |
| HSV_S | $s$ | |
| HSV_V | $v$ | |
| HSV_HdivS | $\frac{h}{s}$ | |
| HSV_HdivV | $\frac{h}{v}$ | |

**Table A3.**
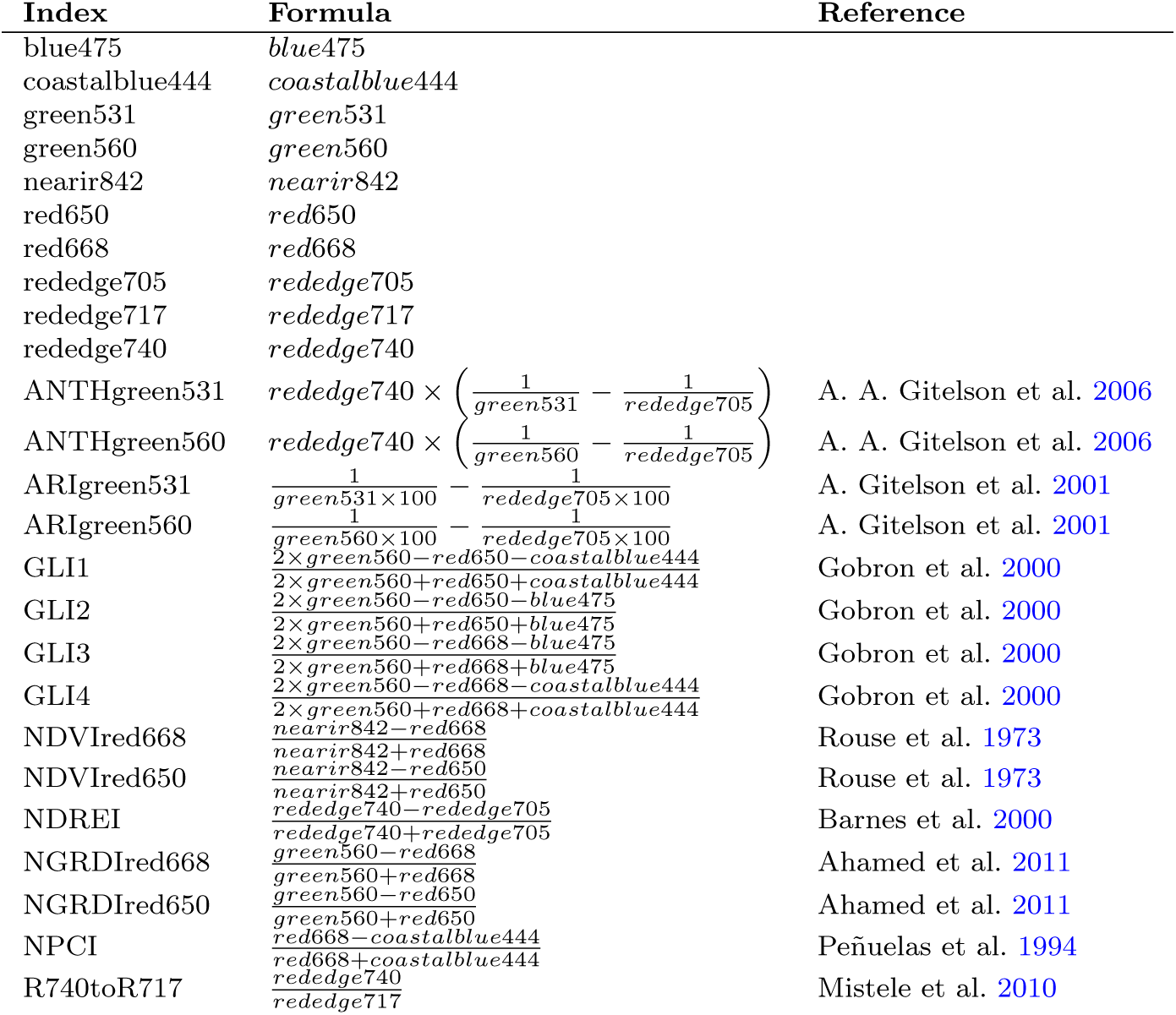
Overview of the applied multispectral indices in this study. The first 10 rows are just each single band from the Micasense sensor.

**Table A4.** Evaluation of the best model using the best RGB (*ExGr*) and SPC (*NDV Ired*650) index. The sum of all correlations for each model was calculated, furthermore the median as well as the range between the minimal and maximal correlation is shown.

| Model | cor sum |  | cor median |  | cor range difference |  |
| --- | --- | --- | --- | --- | --- | --- |
|  | SPC | RGB | SPC | RGB | SPC | RGB |
| C-Gompertz | 4.849 | 4.165 | 0.718 | 0.678 | 0.225 | 0.898 |
| F-Gompertz | 4.504 | 4.427 | 0.681 | 0.771 | 0.454 | 1.095 |
| P-Splines | 2.875 | 2.002 | 0.737 | 0.66 | 0.229 | 1.128 |
| linear | 2.935 | 1.959 | 0.73 | 0.64 | 0.136 | 1.101 |

**Table A5.** The table shows the correlation and RMSE values calculated between RGB (*ExGR*) and SPC (*NDV Ired*650) measurements. Each parameter was compared per measurement year (2021 and 2022) to show year effects.

| parameter | 2021 |  | 2022 |  |
| --- | --- | --- | --- | --- |
|  | correlation | RMSE | correlation | RMSE |
| M | 0.52 | 16.633 | 0.729 | 5.397 |
| onsen | 0.114 | 5.527 | 0.658 | 6.169 |
| endsen |  |  | 0.536 | 5.144 |
| t80 | -0.002 | 7.997 | 0.552 | 6.76 |
| t50 | 0.4 | 11.863 | 0.725 | 5.633 |
| t20 | 0.307 | 21.06 | 0.667 | 5.2 |
| Integral | 0.231 | 92.098 | 0.718 | 56.96 |

**Table A6.**
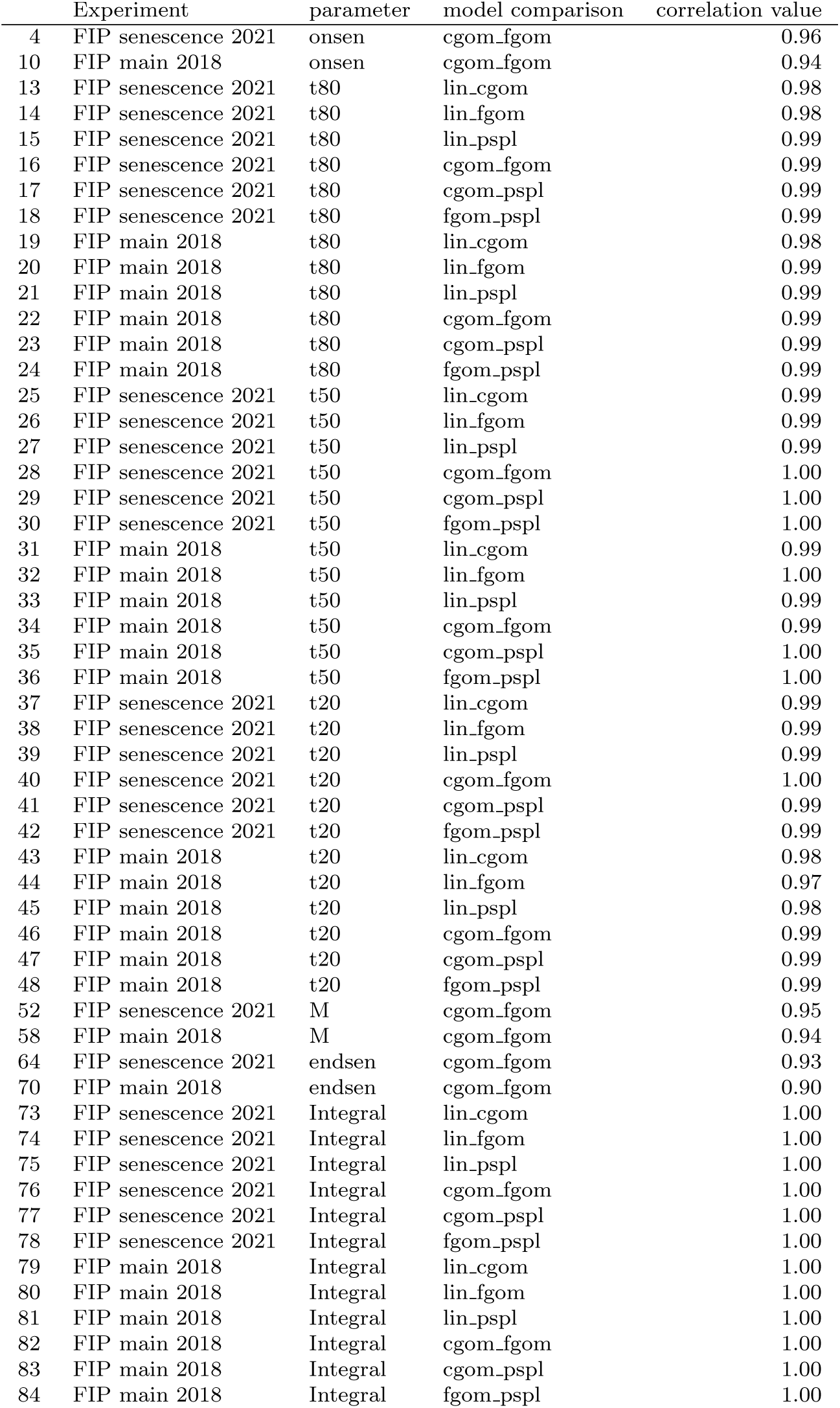
The table shows the correlation between the different models applied to the visual scoring done in two different experiments (FIP senescence 2021 and FIP main 2018). See Figure A8 for parameter *t*80 in the experiment of 2021 as example.

|  | Experiment | parameter | model comparison | correlation value |
| --- | --- | --- | --- | --- |
| 4 | FIP senescence 2021 | onsen | cgom_fgom | 0.96 |
| 10 | FIP main 2018 | onsen | cgom_fgom | 0.94 |
| 13 | FIP senescence 2021 | t80 | lin_cgom | 0.98 |
| 14 | FIP senescence 2021 | t80 | lin_fgom | 0.98 |
| 15 | FIP senescence 2021 | t80 | lin_pspl | 0.99 |
| 16 | FIP senescence 2021 | t80 | cgom_fgom | 0.99 |
| 17 | FIP senescence 2021 | t80 | cgom_pspl | 0.99 |
| 18 | FIP senescence 2021 | t80 | fgom_pspl | 0.99 |
| 19 | FIP main 2018 | t80 | lin_cgom | 0.98 |
| 20 | FIP main 2018 | t80 | lin_fgom | 0.99 |
| 21 | FIP main 2018 | t80 | lin_pspl | 0.99 |
| 22 | FIP main 2018 | t80 | cgom_fgom | 0.99 |
| 23 | FIP main 2018 | t80 | cgom_pspl | 0.99 |
| 24 | FIP main 2018 | t80 | fgom_pspl | 0.99 |
| 25 | FIP senescence 2021 | t50 | lin_cgom | 0.99 |
| 26 | FIP senescence 2021 | t50 | lin_fgom | 0.99 |
| 27 | FIP senescence 2021 | t50 | lin_pspl | 0.99 |
| 28 | FIP senescence 2021 | t50 | cgom_fgom | 1.00 |
| 29 | FIP senescence 2021 | t50 | cgom_pspl | 1.00 |
| 30 | FIP senescence 2021 | t50 | fgom_pspl | 1.00 |
| 31 | FIP main 2018 | t50 | lin_cgom | 0.99 |
| 32 | FIP main 2018 | t50 | lin_fgom | 1.00 |
| 33 | FIP main 2018 | t50 | lin_pspl | 0.99 |
| 34 | FIP main 2018 | t50 | cgom_fgom | 0.99 |
| 35 | FIP main 2018 | t50 | cgom_pspl | 1.00 |
| 36 | FIP main 2018 | t50 | fgom_pspl | 1.00 |
| 37 | FIP senescence 2021 | t20 | lin_cgom | 0.99 |
| 38 | FIP senescence 2021 | t20 | lin_fgom | 0.99 |
| 39 | FIP senescence 2021 | t20 | lin_pspl | 0.99 |
| 40 | FIP senescence 2021 | t20 | cgom_fgom | 1.00 |
| 41 | FIP senescence 2021 | t20 | cgom_pspl | 0.99 |
| 42 | FIP senescence 2021 | t20 | fgom_pspl | 0.99 |
| 43 | FIP main 2018 | t20 | lin_cgom | 0.98 |
| 44 | FIP main 2018 | t20 | lin_fgom | 0.97 |
| 45 | FIP main 2018 | t20 | lin_pspl | 0.98 |
| 46 | FIP main 2018 | t20 | cgom_fgom | 0.99 |
| 47 | FIP main 2018 | t20 | cgom_pspl | 0.99 |
| 48 | FIP main 2018 | t20 | fgom_pspl | 0.99 |
| 52 | FIP senescence 2021 | M | cgom_fgom | 0.95 |
| 58 | FIP main 2018 | M | cgom_fgom | 0.94 |
| 64 | FIP senescence 2021 | endsen | cgom_fgom | 0.93 |
| 70 | FIP main 2018 | endsen | cgom_fgom | 0.90 |
| 73 | FIP senescence 2021 | Integral | lin_cgom | 1.00 |
| 74 | FIP senescence 2021 | Integral | lin_fgom | 1.00 |
| 75 | FIP senescence 2021 | Integral | lin_pspl | 1.00 |
| 76 | FIP senescence 2021 | Integral | cgom_fgom | 1.00 |
| 77 | FIP senescence 2021 | Integral | cgom_pspl | 1.00 |
| 78 | FIP senescence 2021 | Integral | fgom_pspl | 1.00 |
| 79 | FIP main 2018 | Integral | lin_cgom | 1.00 |
| 80 | FIP main 2018 | Integral | lin_fgom | 1.00 |
| 81 | FIP main 2018 | Integral | lin_pspl | 1.00 |
| 82 | FIP main 2018 | Integral | cgom_fgom | 1.00 |
| 83 | FIP main 2018 | Integral | cgom_pspl | 1.00 |
| 84 | FIP main 2018 | Integral | fgom_pspl | 1.00 |

### A.2 Supplementary Figures

**Fig. A1.**
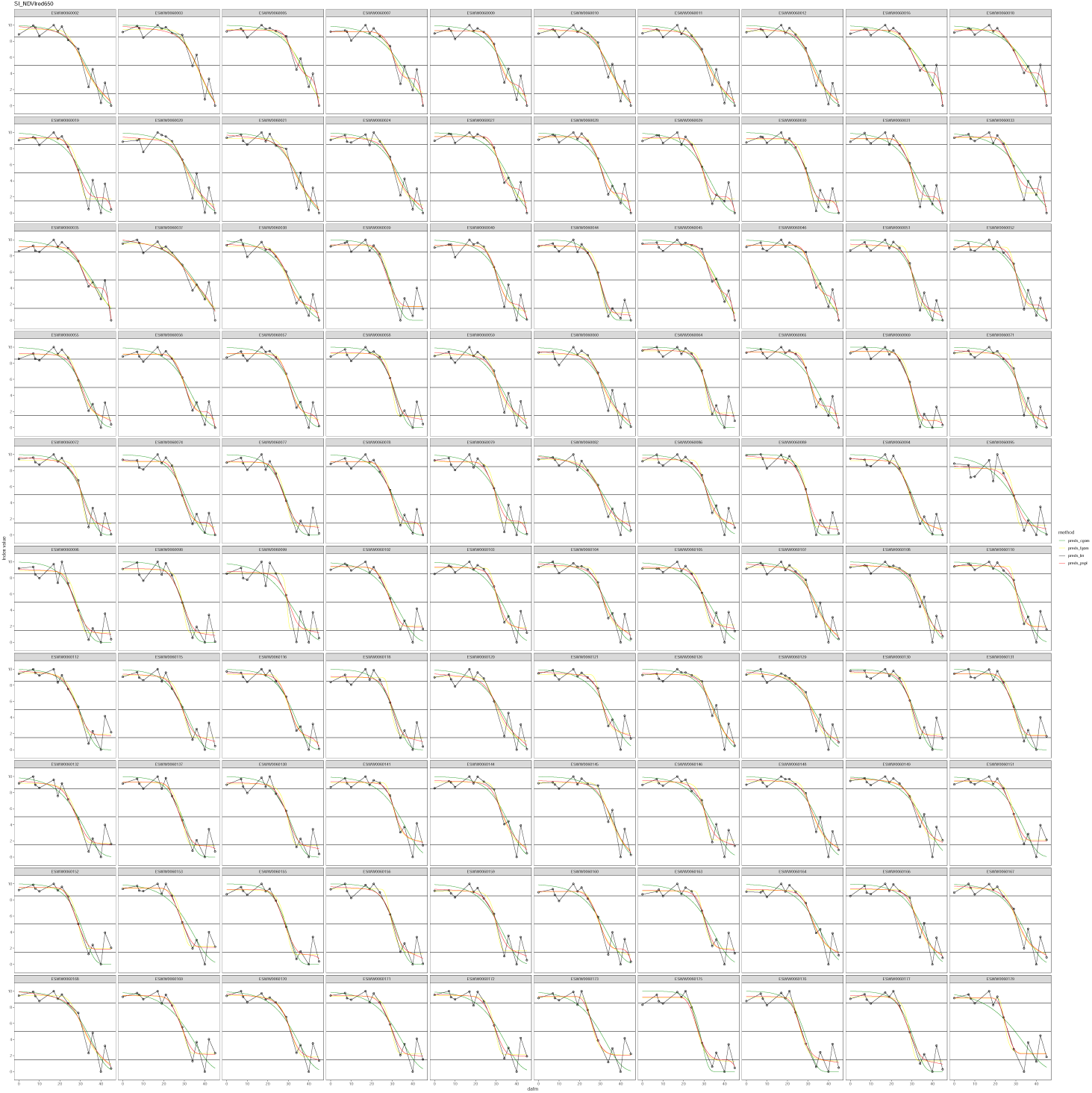
100 random selected plots showing the four different models (lines) as well as the visual scoring (points; y-axis normalized senescence score) over time (x-axis) for the *NDV Ired*650 SPC index.

**Fig. A2.**
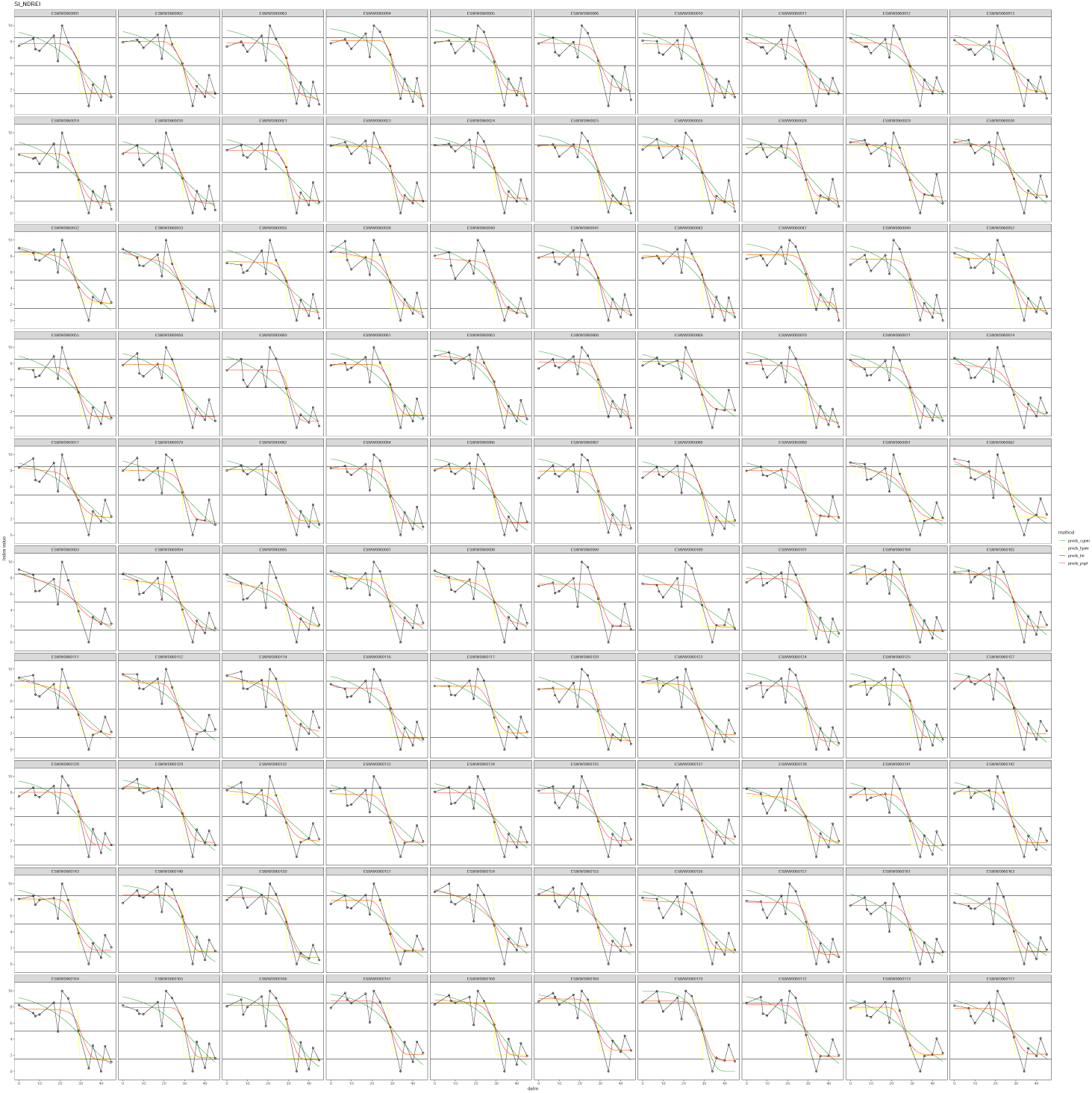
100 random selected plots showing the four different models (lines) as well as the visual scoring (points; y-axis normalized senescence score) over time (x-axis) for the *NDREI* SPC index.

**Fig. A3.**
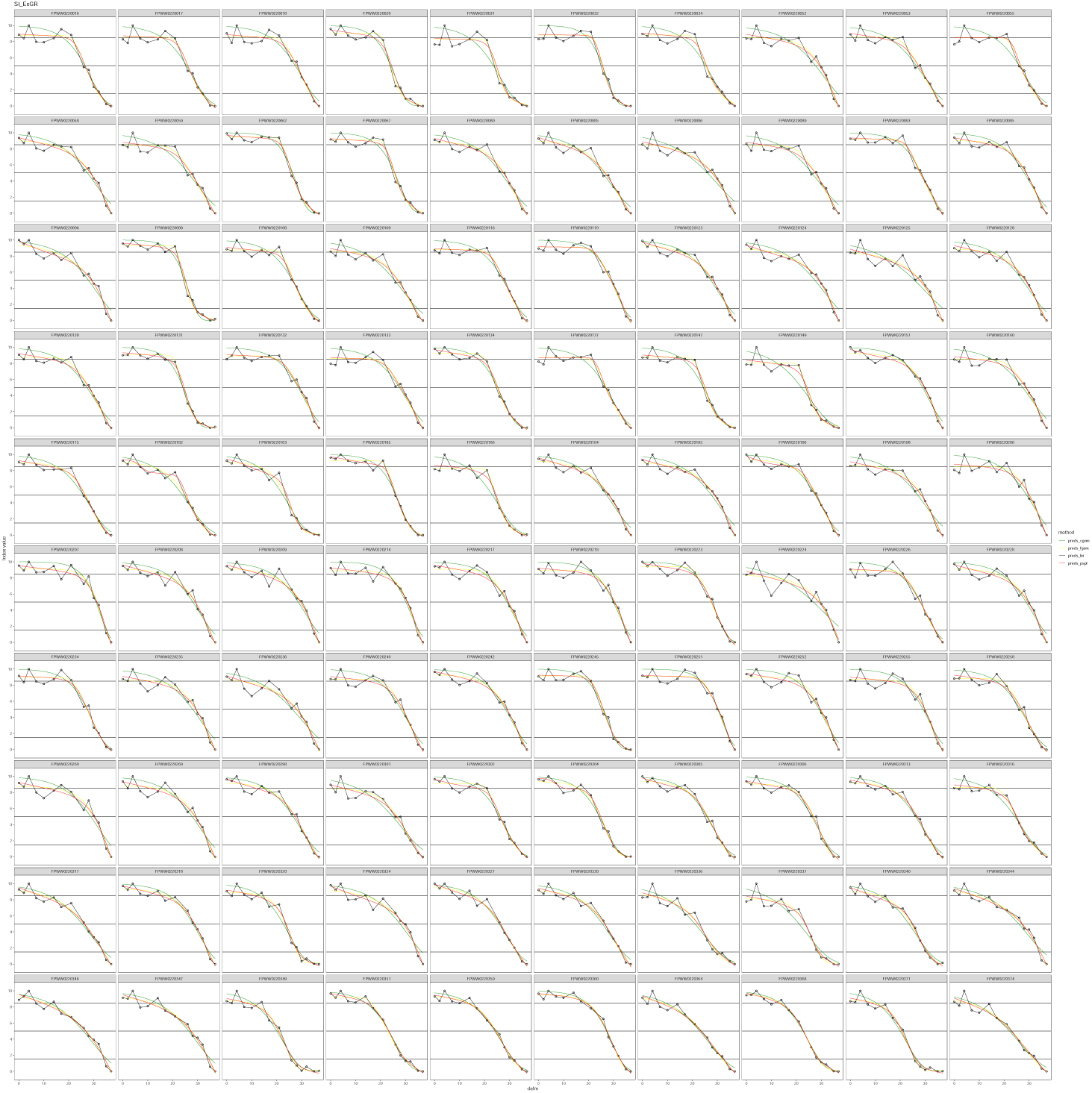
100 random selected plots showing the four different models (lines) as well as the visual scoring (points; y-axis normalized senescence score) over time (x-axis) for the *ExGR* RGB index.

**Fig. A4.**
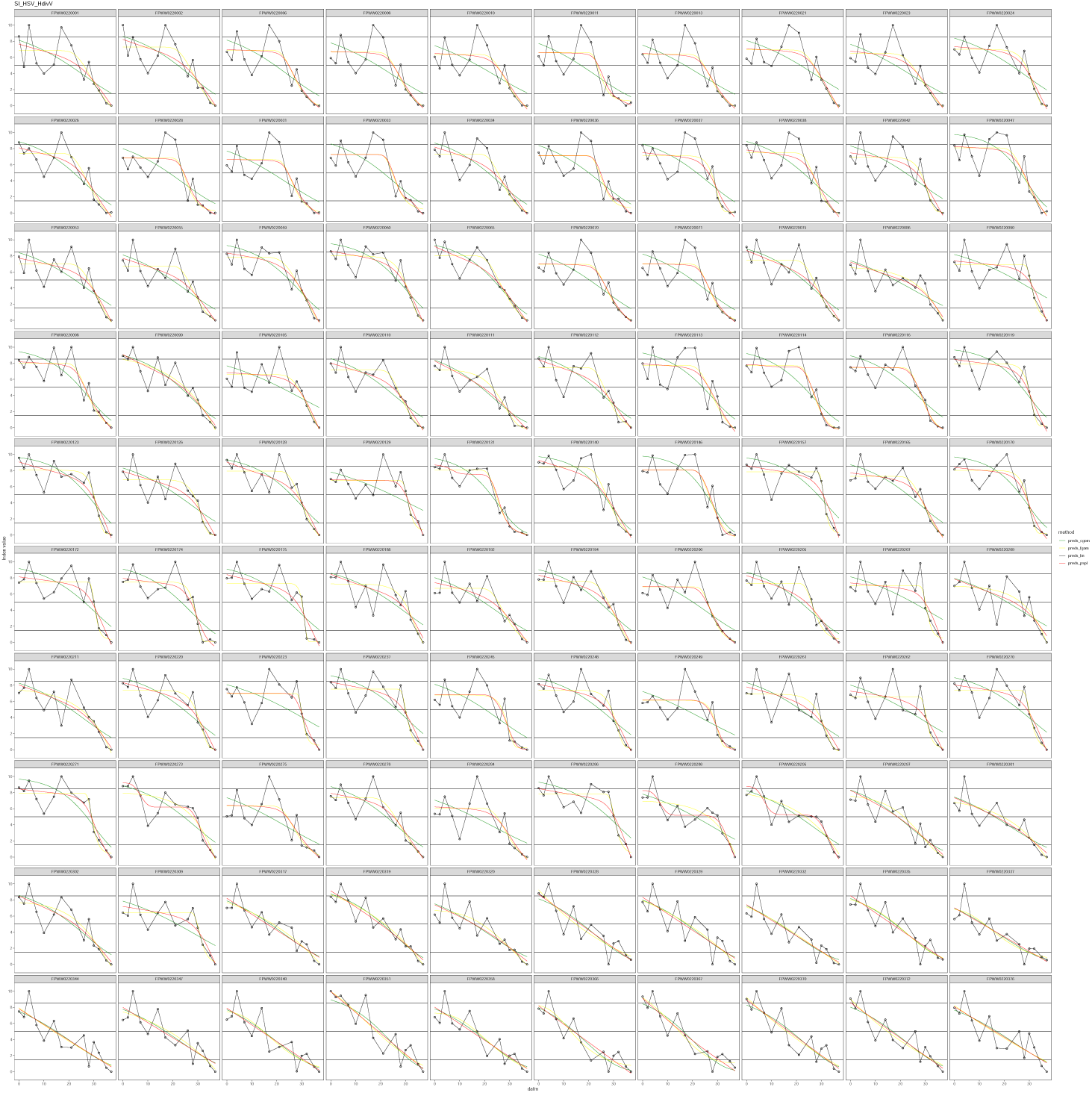
100 random selected plots showing the four different models (lines) as well as the visual scoring (points; y-axis normalized senescence score) over time (x-axis) for the *HSV − HdivV* RGB index.

**Fig. A5.**
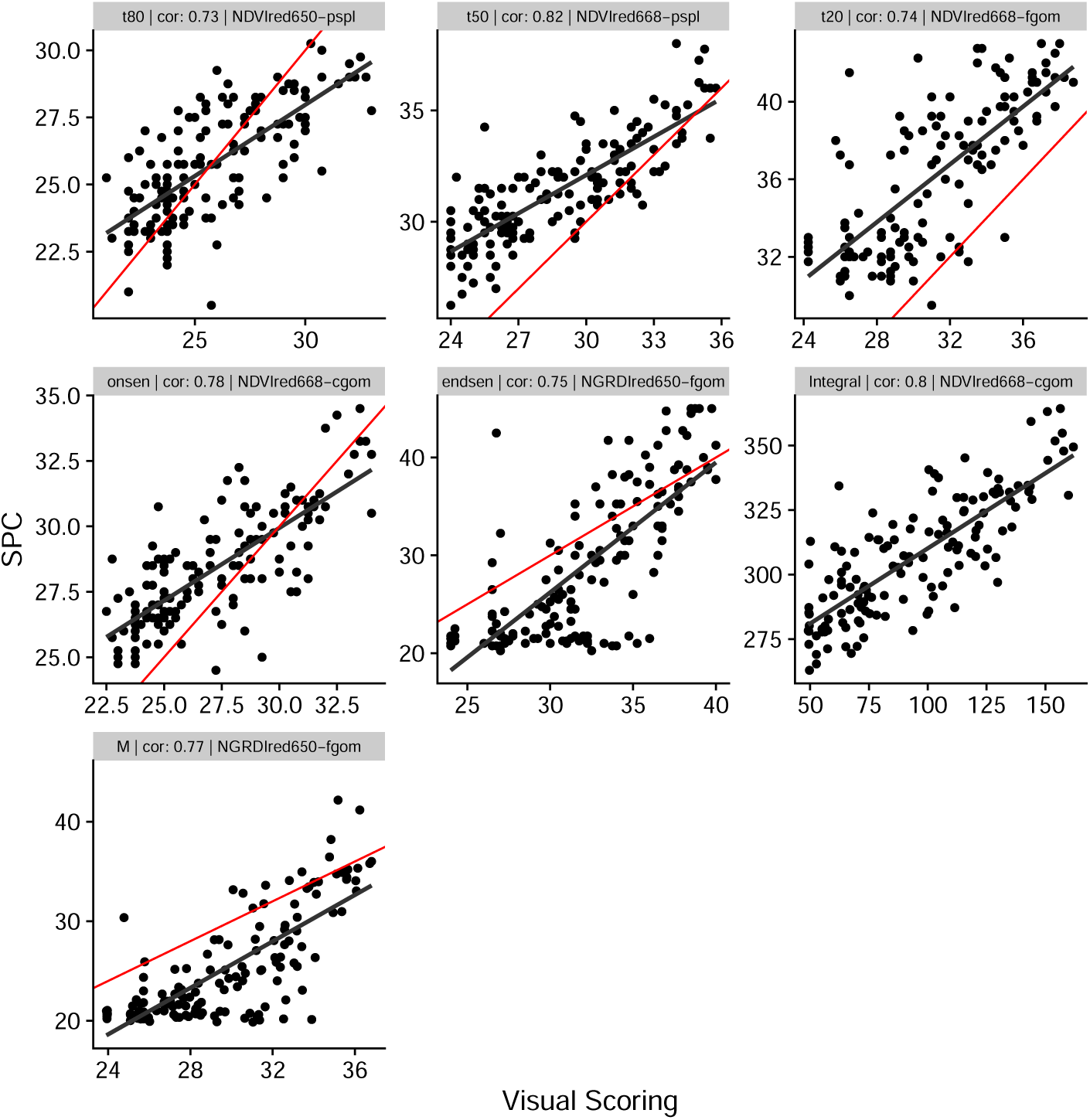
The Figure shows the correlation between the visual scoring (x-axis) and the best SPC model-index combination per parameter measured by high throughput field phenotyping (HTFP). The red line represent the 1:1 line, whereas the black line shows the correlation fit. Each panel shows one parameter (indicated in the header) and the corresponding Pearson correlation value.

**Fig. A6.**
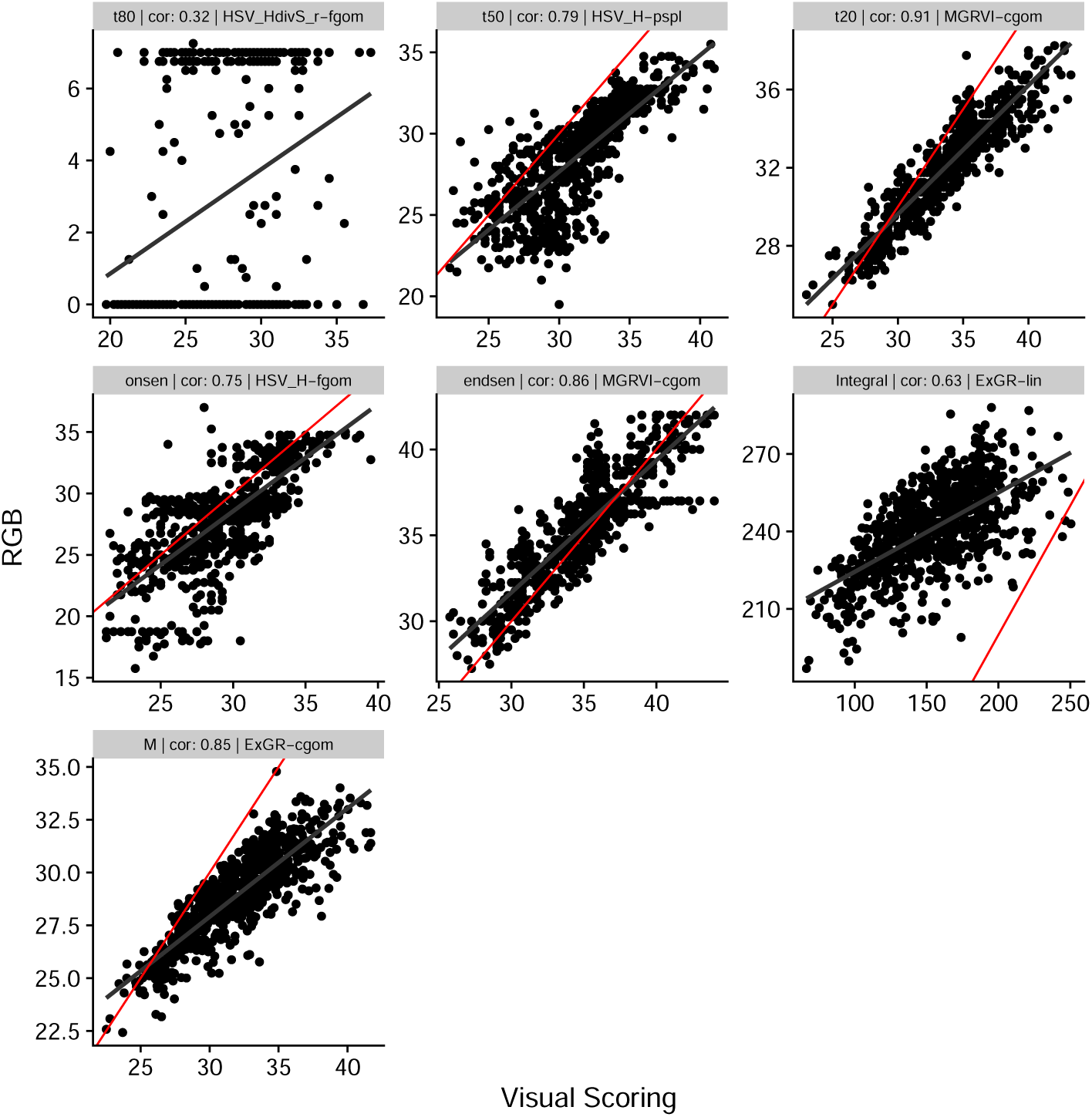
The Figure shows the correlation between the visual scoring (x-axis) and the the best RGB model-index combination per parameter measured by high throughput field phenotyping (HTFP). The red line represent the 1:1 line, whereas the black line shows the correlation fit. Each panel shows one parameter (indicated in the header) and the corresponding Pearson correlation value.

**Fig. A7.**
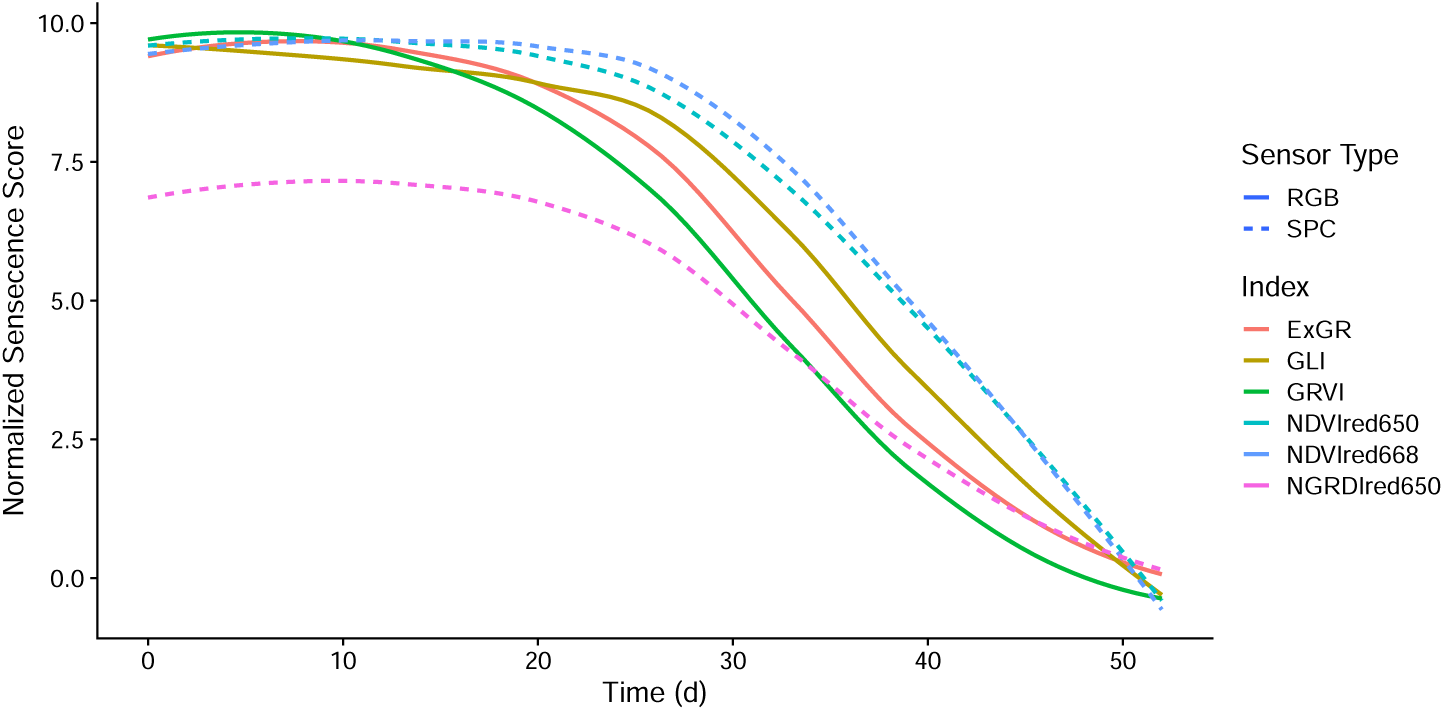
The figure shows the time (in days) on the x-axis and the normalized senescence score values of different indices on the y-axis. For each of the sensors (SPC in dashed lines and RGB in solid lines), three indices were selected, of which the median values over all plots in the FIP main experiment in the year 2022 were calculated and shown here.

**Fig. A8.**
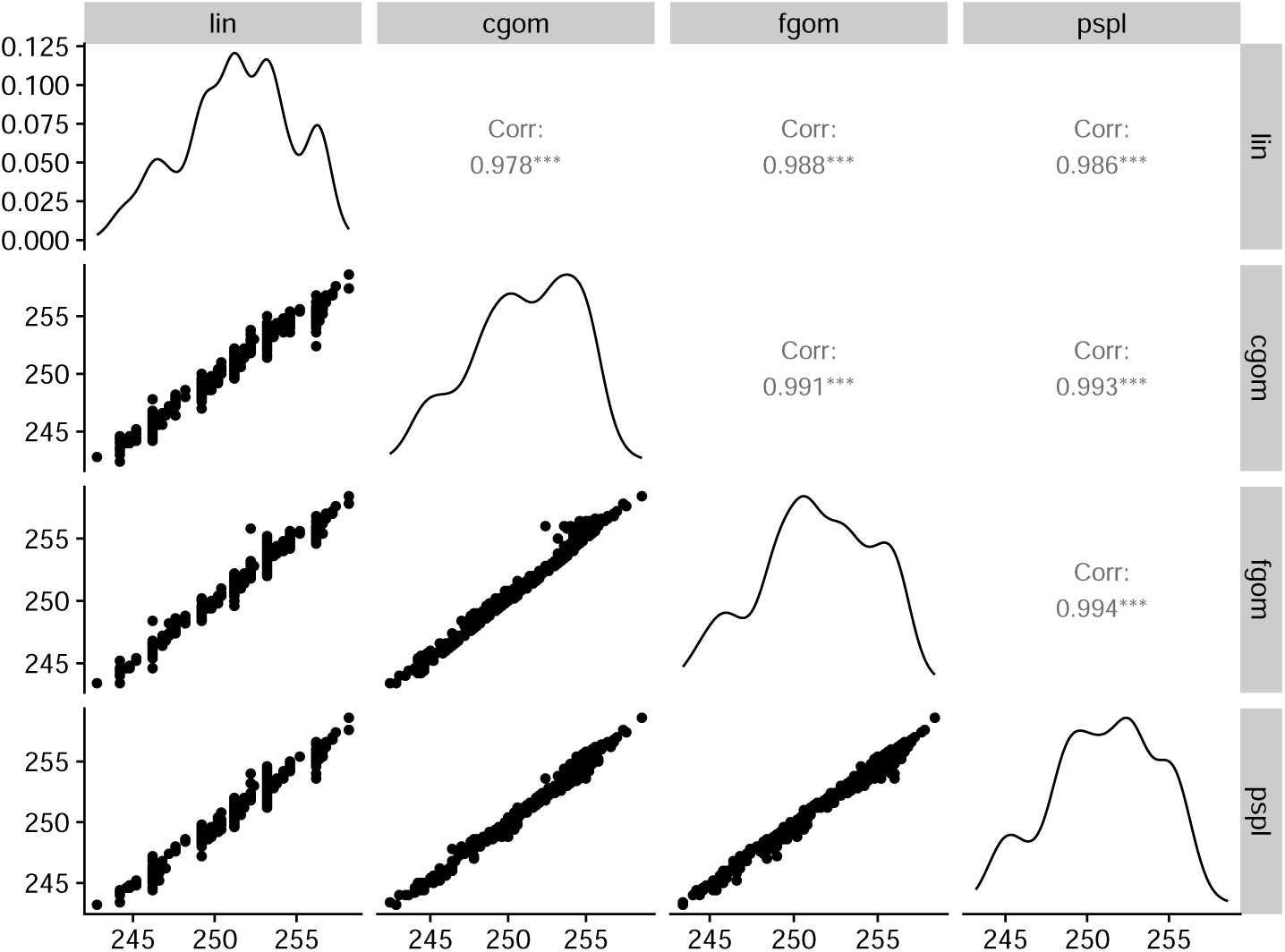
The figure shows the correlation between the different used models applied to the visual scoring data on both of the experiments in which data was available for the parameter *t*80. Each model was correlated crosswise (lower left) and a correlation coefficient was calculated and shown in the correspond panels (upper right). In the middle line the distribution of the data is shown for each model. Comparable correlations are observed for the other parameters.

## Notes

### Competing Interest Statement

The authors have declared no competing interest.

